# Early immune responses anticipate HIV rebound and precede viral control

**DOI:** 10.64898/2026.03.23.713704

**Authors:** Anna Farrell-Sherman, Walker Azam, Natalia de la Force, Ellen L. White, Demi A. Sandel, Antonio E. Rodriguez, Tony Figueroa, Thomas Dalhuisen, Meghann C. Williams, Viva Tai, George W. Gruenhagen, Alaa Abdellatif, Gabriela K. Fragiadakis, Peyman Samghabadi, Carlo De la Sancha Verduzco, Poonam Vohra, Cynthia Gasper, Steven Long, Marina Caskey, Bokai Zhu, Alex K. Shalek, Rebecca Hoh, Rachel L. Rutishauser, Michael J. Peluso, Steven G. Deeks, Lillian B. Cohn

**Affiliations:** Vaccine and Infectious Disease Division, Fred Hutchinson Cancer Center, Seattle, WA, United States; Molecular and Cellular Biology Program, University of Washington, Seattle, WA, United States; Department of Medicine, University of California, San Francisco, San Francisco, CA, United States; Department of Pathology, University of California, San Francisco, San Francisco, CA, United States; Laboratory of Molecular Immunology, The Rockefeller University, New York, NY, United States; Ragon Institute of MGH, MIT, and Harvard, Cambridge, MA, United States; Broad Institute of MIT and Harvard, Cambridge, MA, United States; Institute of Medical Engineering and Science, Massachusetts Institute of Technology, Cambridge, MA, United States; Department of Chemistry, Massachusetts Institute of Technology, Cambridge, MA, United States; Koch Institute for Integrative Cancer Research, Massachusetts Institute of Technology, Cambridge, MA, USA; Department of Global Health, University of Washington, Seattle, WA, United States

**Author notes:** These authors contributed equally to the work (Anna Farrell-Sherman, Walker Azam, Natalia de la Force). These authors jointly supervised the work (Steven G. Deeks, Lillian B. Cohn).

## Abstract

Sustained viral suppression following antiretroviral treatment (ART) cessation is a major goal of HIV cure research^1^. Rare individuals mount immune responses able to control viral rebound without intervention^2,3^, however, the earliest moments in which these responses form remain poorly defined. We performed an intensively sampled, prospective analytical treatment interruption (ATI) to study the initial immune response to rebound and to understand its role in defining subsequent virus control. Profiling of peripheral blood mononuclear cells and plasma revealed consistent immune activation prior to systemic rebound, including upregulation of antiviral transcriptional pathways, expansion of CD16^++^ non-classical monocytes, and increases of inflammatory and antiviral soluble plasma proteins. Individuals with prior viral control (controllers) diverged from non-controllers with a slower slope of rebound, a longer period of immune activity prior to rebound, and engagement of a multifaceted immune program with less systemic inflammation. An intermediate immune signature emerged in a separate ATI cohort of individuals who experienced delayed rebound after receiving broadly neutralizing antibodies^4^, suggesting that immunotherapy can induce a potentially protective pre-rebound immune response. Together, these data resolve the earliest systemic host immune responses to HIV rebound and demonstrate broad immune differences associated with HIV control phenotypes.

## INTRODUCTION

Antiretroviral therapy (ART) has transformed HIV into a manageable chronic disease^5^. However, ongoing variability in HIV/AIDS services worldwide highlights the vulnerability of care systems for people living with HIV (PLWH), and has added urgency to the global effort to develop a safe, effective, and scalable cure strategy^1,6^. While consistent ART suppresses plasma viremia in most PLWH^7–9^, it is not curative because the proviral reservoir persists and can seed rebound after ART is interrupted^10–12^. In most cases, virus becomes detectable 2 to 3 weeks after treatment interruption, expands exponentially, then returns to a set-point similar to pre-ART chronic viral loads^13^. Most PLWH have a set-point above 10,000 copies/ml, but a small subset can achieve sustained partial control (viral load <2,000 copies/ml, viremic controllers) ^2,14^. Importantly, people who control HIV prior to ART typically also control the virus after ART cessation^15^, and pre-ART viral load is a predictor of time-to-rebound^16^. A major goal of cure-focused research is to design interventions that enable non-controllers to generate controller-like phenotypes, an outcome referred to as post-intervention control or durable ART-free viral suppression^17^.

The host responses associated with viral control at set-point have been extensively studied^17–24^, however mechanistic insight into the pre-set-point events contributing to viral control is limited. Studies show that: immune activation can be detected before viremia becomes systemic^25–27^; early viral growth rates differ between control phenotypes^28–30^; and, immune modulating interventions can affect rebound dynamics^30–32^. Rebound likely begins with very low-level viral replication in tissues^33–35^ with multiple initial “intercepts” between the host immune system and newly activated virus that occur well before viremia becomes detectable in the blood. We hypothesize that the immune responses at these intercepts are critical for mounting systemic immune responses that shape the rebound trajectory. Unfortunately, this intercept period is challenging to study. In most analytical treatment interruption (ATI) studies, sampling is too infrequent to disentangle immune changes at initial rebound from those that reflect a response to established viremia. This distinction is key for designing interventions that allow more individuals to become post-intervention controllers.

Here, we examine the immune response during the host-virus intercept when the virus first re-emerges and encounters host immunity. We conducted a densely sampled, prospective ATI study (“Researching Early Biomarkers of Unsuppressed HIV Dynamics,” REBOUND). REBOUND specifically studied responses in the absence of immune-modulating interventions, sampled at high temporal resolution during ATI, and enrolled individuals with diverse pre-ART viral control patterns. Using transcriptomics, cell subset analysis, and plasma proteomics, we interrogated the host response to rebound during the host-virus intercept. We show that coordinated innate activation precedes rebound and that controllers engage broader functional immune programs earlier than non-controllers. Our results allow us to dissect the mechanisms that allow for viral control *in vivo* and could inform the immunological targets of future interventional cure studies.

## RESULTS

### The REBOUND study interrogates early immune response to HIV-1 rebound viremia

The REBOUND study enrolled twenty PLWH into an intensively sampled observational ATI (NCT04359186, **Fig 1a**). Peripheral blood mononuclear cell (PBMC) and plasma samples were collected up to three times per week following treatment interruption through ART restart. In a subset of participants, lymph node tissue fine needle aspirates and leukaphereses were collected at baseline on ART and during the ATI, timed to coincide with predicted intercept (8-24 days into the ATI).

**Fig. 1.**
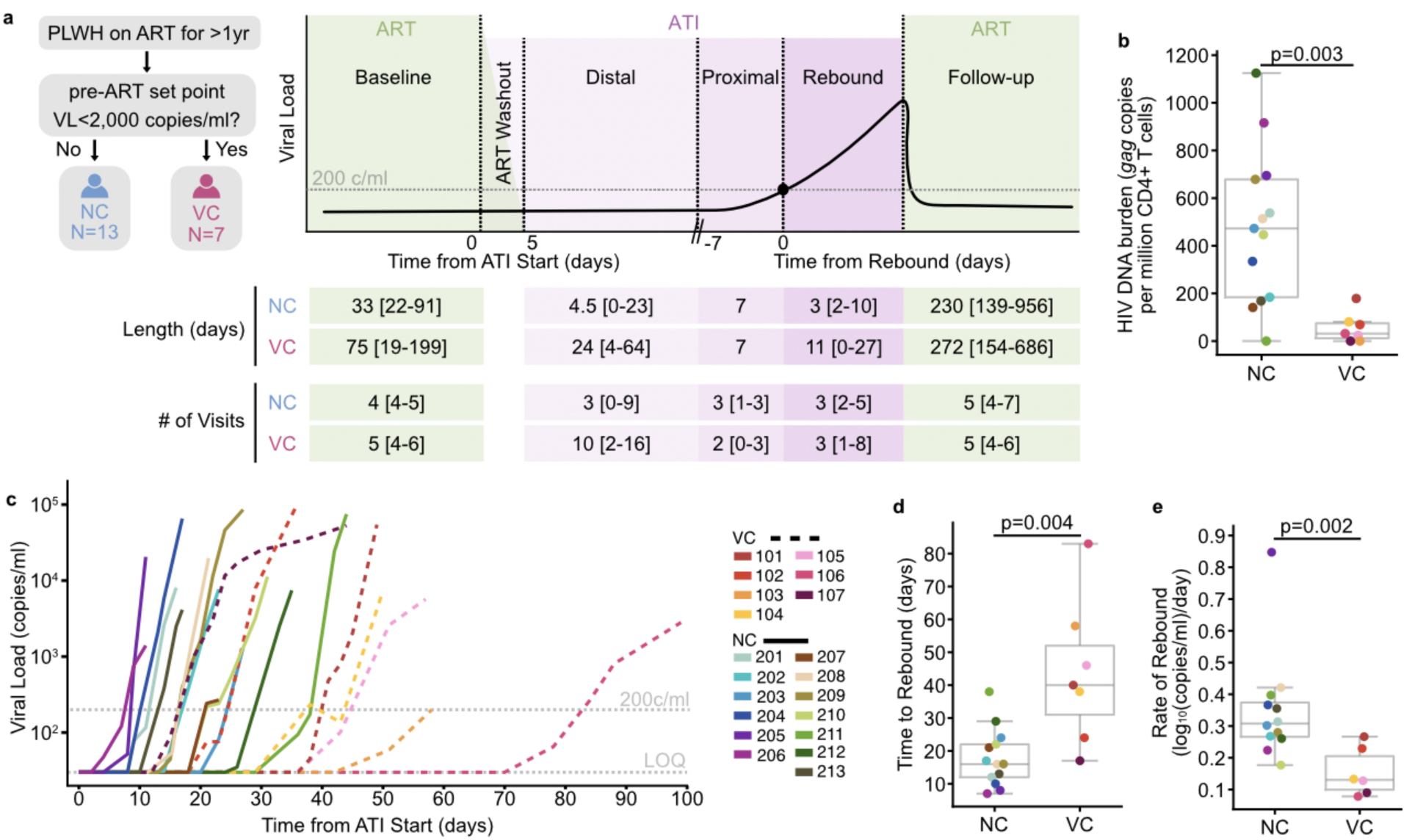
The REBOUND study: an observational analytical treatment interruption (ATI) with intensive sampling. **a**, REBOUND study schematic defining controller status classification and study sampling stages: baseline (pre-ATI timepoints on ART); ART washout (the pharmacological tail of ART, 1-4 days after ATI start); distal (day 5 after ATI start until proximal); proximal (7 days preceding rebound); rebound (viremic timepoints after estimated date of rebound); follow-up (post ART reinitiation). Rebound category length and number of visits per category is displayed with median and range per participant in each controller phenotype category. **b**, HIV proviral DNA burden per participant as measured by *gag* copies per million CD4^+^ cells for viremic controllers and non-controllers. **c,** Viral load curves of participants; viremic controllers (VC, dashed lines) and non-controllers (NC, solid lines). Limit of quantification (LOQ) for viral load was 30 copies/ml. **d**, Time to rebound and **e**, rate of rebound between VC and NC. All p-values from two-sided Mann-Whitney U tests.

Since low viral set-point in viremic controllers (VC) is re-achieved after ART interruption^36^, we used historical pre-ART viral load measurements and clinical documentation to assign controller status at study enrollment. Seven participants met criteria for viremic control, defined by having a median pre-ART plasma viral load (VL) <2,000 copies/ml (ART started at a median of 4.8 years after estimated HIV acquisition, range 1.7-23.6). The remaining 13 participants with no prior evidence of viral control were classified as non-controllers (NC); 12 initiated ART during chronic infection (median 5 years after estimated HIV acquisition; range 0.92-16.1), and one initiated ART 7 days after HIV acquisition and had no detectable HIV-specific cellular or humoral immunity^37^. Individual participant viral load curves are shown in **Extended Data Fig. 1-2**; ART restart criteria are included in the **Methods**.

To assess reservoir burden at baseline, we quantified proviral *gag* in CD4^+^ T cells. Proviral *gag* levels were significantly higher in NCs compared to VCs (p=0.003, **Fig. 1b**). VCs and NCs did not differ across other key clinical measures including the number of protective HLA-I alleles, time untreated before ART, or CD4^+^ T cell nadir, although time on ART was longer in NCs than VCs (18.9 years versus 8.2 years, p=0.01, two-sided Mann-Whitney U test). Participant characteristics shown in **Extended Data Table 1**.

The ATI was safe for all participants with no serious adverse events and all participants resuppressed viremia upon ART reinitiation.

### Controllers rebound later and slower than non-controllers

Rebound dynamics varied across participants and corresponded to their pre-ART control phenotype (**Fig. 1c**). We set the threshold for defining rebound at plasma VL ≥200 copies/ml, consistent with the clinical definitions of viral suppression on ART^38,39^, then estimated the date of rebound for each participant (**Methods**) and observed different median times to rebound of 17 days for NCs and 40 days for VCs (p=0.004, **Fig. 1d**). The rate of rebound also differed, with steeper rebound in NC than VC (+0.31 versus +0.13; p=0.002, **Fig. 1e**). Controller status was correlated with days to rebound and negatively correlated with reservoir burden and rebound slope (**Extended Data Fig. 3**), as has been observed in prior studies^29^. Whereas earlier rebound in NCs may reflect higher reservoir burden, the shallower rebound slope in VCs suggests partial immune containment of emerging viremia during the earliest stages of rebound.

### Elevated baseline immune activation in non-controllers on ART

To test whether pre-ATI immune differences could contribute to divergent rebound dynamics, we profiled baseline immune features across participants using repeated sampling of transcriptomics, plasma proteomics, and immune cell phenotyping. Baseline PBMC gene expression differed between VCs and NCs (**Fig. 2a**, **Supplementary Table 1**), with NCs showing higher enrichment of inflammatory and immune activation signals, whereas VCs were enriched for cell cycle-associated signatures (**Fig. 2b**, **Supplementary Table 2**). Lymph tissue, which is assumed to be a possible origin of HIV reactivation after treatment cessation^33–35^, had concordant transcriptional patterns; single-cell RNA-seq of lymph node mononuclear cells from a subset of participants (NC=9, VC=3) showed greater inflammatory pathway enrichment in NC across multiple cell types (**Fig. 2c**, **Supplementary Table 3**). No circulating plasma proteins significantly differed between the groups at baseline (**Supplementary Table 4**). NCs had a higher frequency of inflammatory classical monocytes (CD14^++^ CD16^-^) in the blood compared to VCs (**Fig. 2d**, p=0.001), while non-classical monocytes (CD14^+^ CD16^++^), which are specifically antiviral in nature, were not significantly different between the two groups (**Fig. 2e**, p=0.087) ^40^. Overall, we observe greater levels of baseline inflammation in NC than VC.

**Fig. 2.**
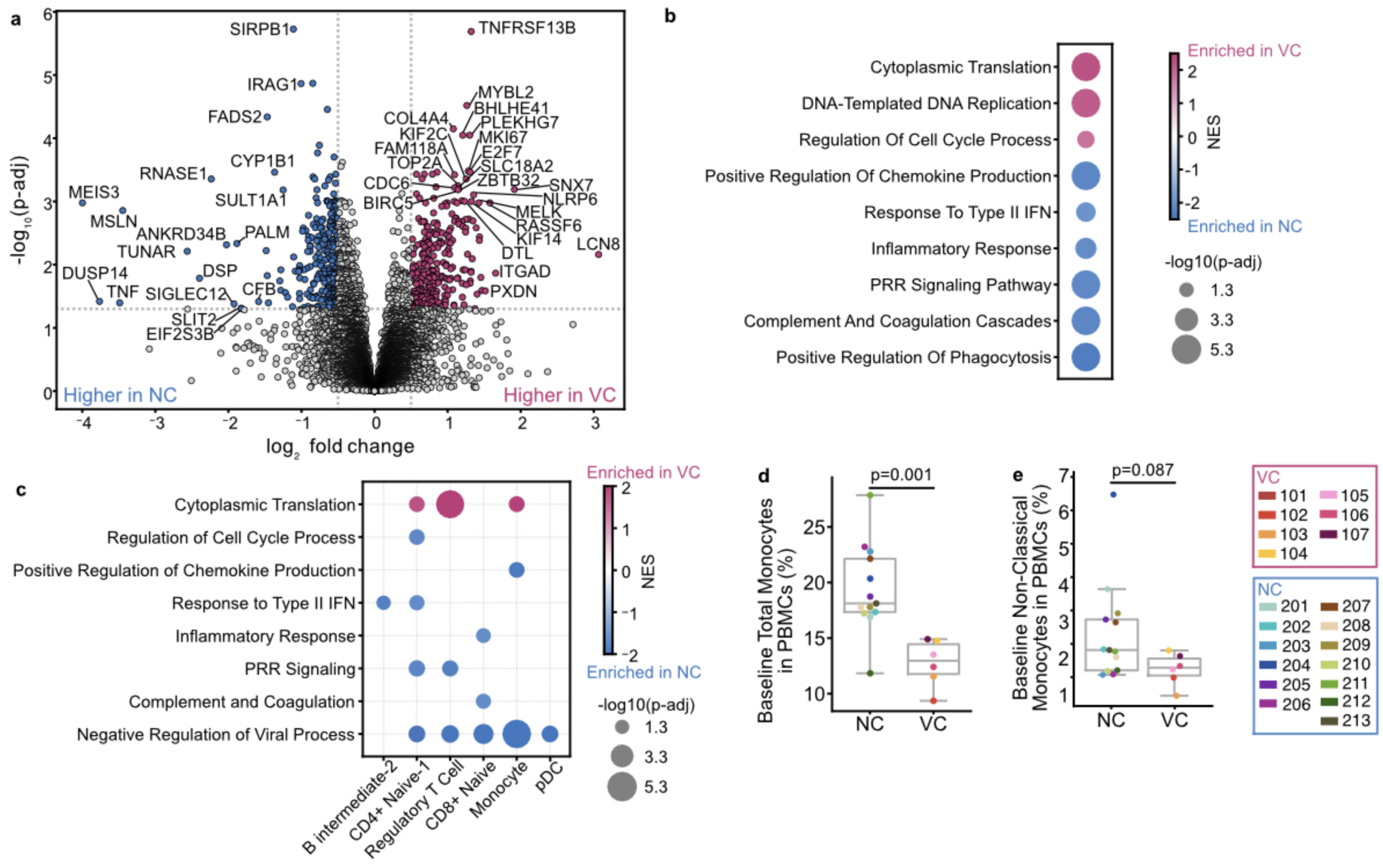
On ART, NC show higher levels of immune activation than VC. **a**, Differential expression of genes between NC (negative) and VC (positive) while on ART. Colored points are significantly differentially expressed genes with |log_2_FC|>0.5 and a p-adj<0.05 (threshold visualized with dotted grey lines). Labeled genes have |log_2_FC|>1.5 and a -log_10_(p-adj)>3. **b**, Differential pathway expression in blood between baseline samples from VC (positive NES) and NC (negative NES). **c**, Differential pathway expression in cell subsets in the baseline lymph nodes of VC (positive NES) and NC (negative NES). Full pathway names and reference numbers in Extended Data Table 2. **d**,**e**, Comparison of total monocyte abundance (**d**) and non-classical monocyte abundance (**e**) as a percentage of PBMCs between VC and NC at baseline. p-values from two-sided Mann-Whitney U test. NES = Normalized Enrichment Score.

### Increases in antiviral proteins and immune cell activation at the intercept

To assess whether antiviral responses emerge during the host-virus intercept, we quantified the abundance of 249 immune plasma proteins across 276 plasma samples collected throughout the ATI (median 14 samples per participant). To assess differences in kinetics, we defined five ATI stages across the study (**Fig. 1a**). Baseline timepoints were collected prior to ART interruption (median of 4 per participant, sampled 0-199 days before ATI start). ATI timepoints prior to rebound are classified as distal (after pharmacologic waning; ATI day ≥5 through >7 days before rebound) or proximal (≤7 days prior to rebound). Rebound timepoints are those sampled on or after the date of rebound. Timepoints sampled after ART re-initiation are considered follow-up (3-184 days after restart).

Individual protein abundances were largely unchanged from baseline to the distal stage and only IFN-⍺1 increased from baseline to proximal (p-adj=0.036, **Fig. 3a**). At the rebound stage, 50 proteins were differentially abundant relative to baseline, including canonical inflammatory and antiviral mediators such as IFN-⍺1, IFN-𝛾, IFN-λ1, IL12p70, IL-15, and CXCL10 (**Fig. 3a**, **Supplementary Table 4**). Throughout pre-rebound and the early growth phase of viremia, IFN-⍺1 tracked with plasma viral load in most participants (**Extended Data Fig. 4**). We also confirmed an increase in soluble LAG3, a known marker of immune activation^41^, which we observed in an earlier cohort^25^, while IL-6, often associated with chronic HIV inflammation^42^, did not change.

**Fig. 3.**
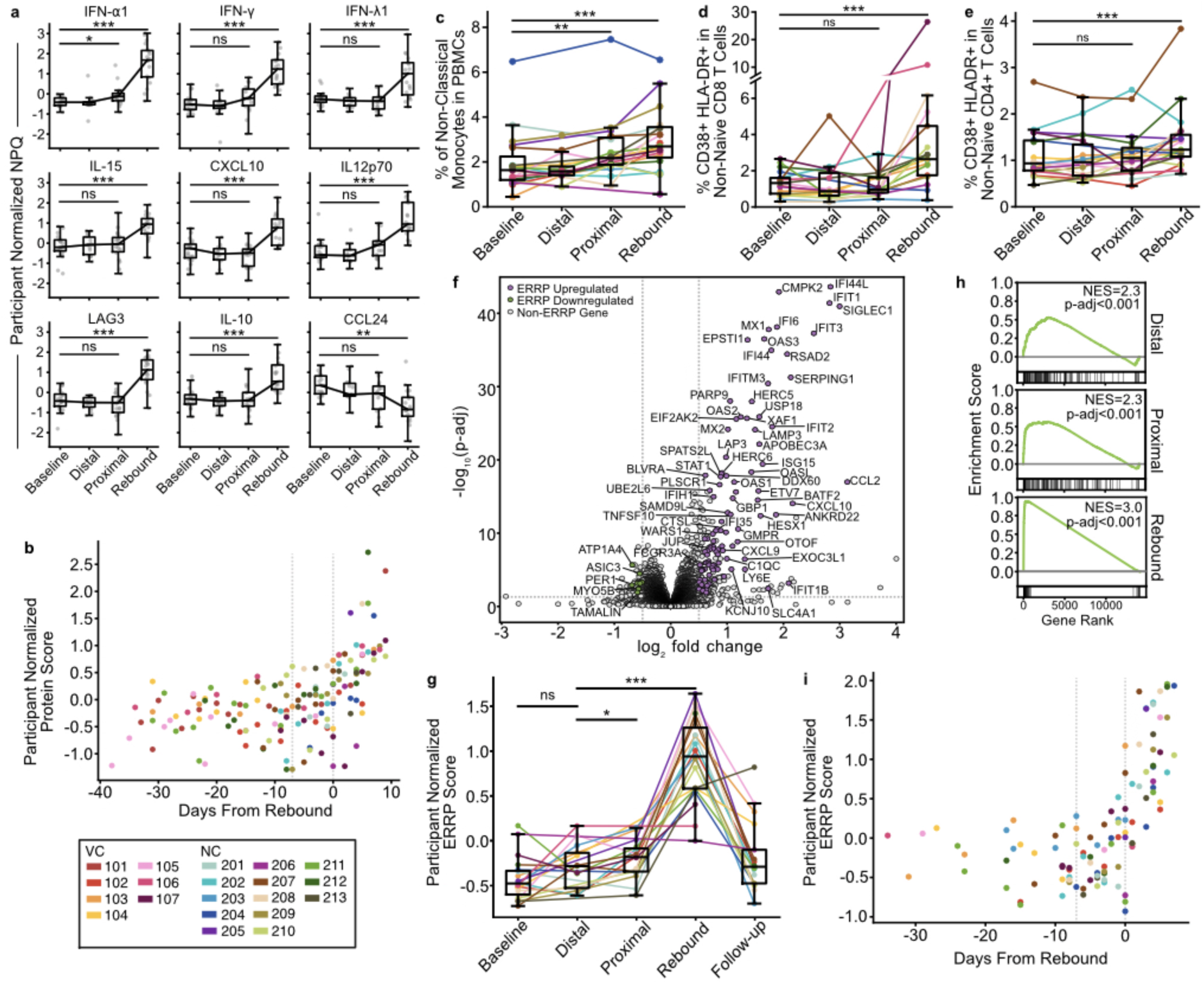
Antiviral immune activation at the intercept. **a**, Plasma protein abundance by NULISA Protein Quantification (NPQ) normalized by participant. FDR-values from two-sided Wilcoxon signed-rank test using median participant value per ATI stage. **b**, Association between immune protein score and days from rebound. Points colored by participant. Vertical dotted lines at the first proximal timepoint (-7 days) and rebound (0 days). **c**,**d**,**e** Proportion of non-classical monocytes in total PBMCs (**c**) and CD38^+^ HLA-DR^+^ cells in non-naïve CD8^+^ T cells (**d**) and CD38^+^ HLA-DR^+^ cells in non-naïve CD4^+^ T cells (**e**) per participant at each ATI stage. p-values from a two-sided paired Wilcoxon signed-rank test between participant medians. **f**, Differential expression of genes between baseline (negative) and rebound (positive). Colored points are Early Rebound Response Pathway (ERRP) genes, with key selected genes labeled. Full list of ERRP genes in Supplemental Table 2. Dotted grey lines represent significance thresholds at |log_2_FC|>0.5 and an p-adj<0.05. **g**, Average participant-normalized EERP gene expression per ATI stage. Points and lines colored by participant. p-values from two-sided Wilcoxon signed-rank test. **h**, ERRP enrichment plot from baseline at distal, proximal, and rebound. **i**, Association between ERRP expression score and days from rebound. Points colored by participant. Vertical dotted lines at the first proximal timepoint (-7 days) and rebound (0 days). All boxplots represent interquartile range and median across participants. * p<0.05, **p≤0.01, *** p≤0.005.

To increase our detection sensitivity to coordinated shifts in protein levels, we aggregated the 50 plasma proteins into a single participant-normalized expression score (**Fig. 3b, Methods**). To assess the relationship between the protein score and days from rebound, we built a continuous participant-specific random-intercept mixed-effects model (**Extended Data Fig. 5**). The score was associated with days from rebound, indicating that a soluble immune protein response emerges before the rebound threshold and scales with proximity to rebound (R^2^_GLMM(c)_=0.49, β=8.21, SE=1.0, p=3.9x10^16^).

As in our previous work^25^, high-dimensional flow cytometry revealed an increase in non-classical monocyte (CD14^+^ CD16^++^) abundance prior to the rebound threshold (baseline to proximal mean FC=1.28, p=0.02, **Fig. 3c**). From baseline to rebound, non-classical monocytes increased further (mean FC=1.77, p=0.0002, **Fig. 3c**), alongside an increase in activated CD4^+^ and CD8^+^ T cells (HLA-DR^+^ CD38^+^; mean FC=2.0, p=0.001, **Fig. 3d-e**). Collectively, these data argue that the intercept period is defined by subtle increases in antiviral immune activation which expand rapidly once virus becomes abundant in blood.

### A new early rebound transcriptional signature emerges at the intercept

To directly assess cellular activity during rebound, we performed bulk transcriptomic profiling of PBMCs and compared baseline to rebound stages using paired differential expression analysis. This analysis identified 116 Differentially Expressed Genes (DEGs) that changed consistently across participants (|log_2_FC|>0.5, p-adj<0.05; **Fig. 3f**; **Supplementary Table 2**). As expected, these DEGs included canonical innate antiviral and interferon stimulated genes (e.g. *IFIT1*, *MX2*, *OAS3*, *CXCL10*) as well as monocyte associated genes, such as CCL2. Several DEGs were less well characterized in this context, including some that have not previously been associated with an immune or antiviral response. We defined these 116 rebound DEGs as the Early Rebound Response Pathway (ERRP).

No individual genes increased significantly prior to the rebound stage, so to identify more subtle expression changes, we assessed ERRP expression as a single score across ATI (**Methods**). The ERRP score increased from the distal to proximal stage (p=0.019), and from distal to rebound (p=0.0002; **Fig. 3g**), but did not change from baseline to the distal stage. Gene Set Enrichment Analysis (GSEA) showed enrichment of ERRP at rebound (p-adj<0.001), distal (p-adj<0.001) and proximal (p-adj<0.001; **Fig. 3h**). ERRP expression returned to baseline levels after ART reinitiation (baseline vs follow-up p=0.056). Finally, in a continuous participant-specific random-intercept mixed-effects model the ERRP score was associated with days from rebound (R^2^_GLMM(c)_=0.51, β=6.62, SE=0.82, p=3.37x10^-15^; **Fig. 3i, Extended Data Fig. 5**). These results indicate that a coordinated transcriptional response is evident soon after ART cessation during the intercept period.

### Earlier and broader immune engagement in controllers at intercept

Having defined the intercept immune response across participants, we next tested the main hypothesis of our study: that early immune responses to rebound vary between individuals with different control phenotypes.

VCs showed earlier engagement of cellular immune responses. VCs exhibited larger pre-rebound expansions of antiviral non-classical monocytes than NCs (baseline to proximal FC=1.6 vs 1.2, p=0.048; baseline to rebound FC=2.2 vs 1.3, p=0.022, **Fig. 4a**). Based on mass cytometry (CyTOF) profiling (five samples per participant), VCs also had higher frequences of activated/cycling non-naïve CD8^+^ T cells at rebound (CD38^+^ HLA-DR^+^ median 3.4% vs 0.76%; p=0.012; Ki-67^+^ median 5.33% vs 1.95%; p=0.007; **Fig. 4b-c**). In contrast, NCs had higher IFN-⍺1 at rebound (p=2.4x10^-4^, **Fig. 4d**), while other interferons did not differ. These differences were not explained by viral loads during rebound, which were similar between groups (p=0.53, **Extended Data Fig. 6**). These results suggest that VCs engage earlier and more functional cellular immune programs with less systemic inflammatory activation than NCs, a pattern consistent with subsequent containment of emerging viremia.

**Fig. 4.**
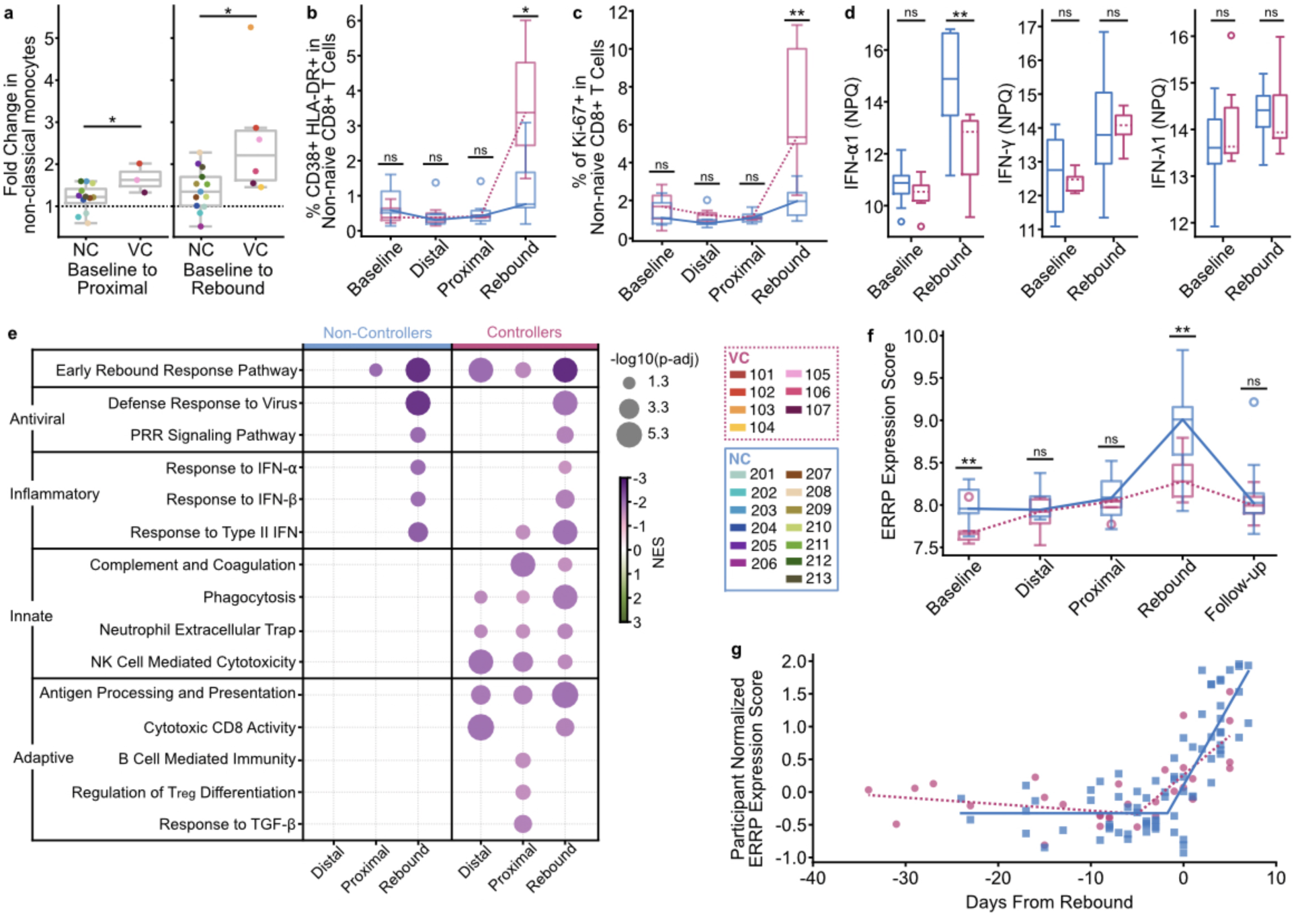
Viremic Controllers show earlier, multi-arm immune engagement at intercept. **a**, Fold change increase in non-classical monocyte abundance from baseline to proximal (left) and baseline to rebound (right). Points colored by participant. p-values from two-sided Mann-Whitney U test. **b**,**c,** Proportion of non-naïve CD8^+^ T cells expressing CD38 and HLA-DR (**b**) or Ki-67 (**c**) in VC (dotted line) and NC (solid line) at each rebound category. p-values from two-sided Mann-Whitney U test between VC and NC. **d**, Plasma protein abundance by NULISA Protein Quantification (NPQ) between NC (left, solid median) and VC (right, dotted median) at baseline and rebound. p-values from two-sided Mann-Whitney U test between NC and VC participant medians. **e**, Pathway enrichment across ATI stage in NC (left) and VC (right). Positive NES indicates enrichment at indicated ATI stage; negative NES indicates enrichment at baseline. Insignificant (p-adj≥0.05) pathways’ circles not displayed. Full pathway names and reference numbers in Extended Data Table 2. **f**, ERRP expression score (average participant normalized expression of ERRP genes) between VC (dotted line) and NC (solid line) at each rebound category. p-values from two-sided independent t-test between VC and NC. **g**, Continuous two-segment piecewise linear regression models predicting ERRP expression score in NC (squares, solid line) and VC (circles, dotted line) over days from rebound. All boxplots represent interquartile range and median across participants. * p<0.05, ** p≤0.01, *** p≤0.005. NES = Normalized Enrichment Score.

We next compared transcriptomic pathways activity between control phenotypes. In VCs, GSEA showed ERRP enrichment from baseline to distal (p-adj<1x10^-5^), proximal (p-adj=0.008), and rebound (p-adj<1x10^-5^) stages. In NCs enrichment from baseline was only significant at proximal (p-adj=0.036) and rebound (p-adj<1x10^-5^) (**Fig. 4e**). ERRP was one of the first inflammatory or antiviral signatures enriched during the intercept. Beyond ERRP, VCs displayed enrichment of a diverse set of effector pathways, including NK cell mediated cytotoxicity, phagocytosis, neutrophil function, cytotoxic T cell responses, antigen processing, and B cell immunity, which did not reach significance in NCs (**Fig. 4e, Extended Data Table 2**, **Supplemental Table 3**). VCs showed enrichment of regulatory T cell differentiation and TGF-β response at the proximal stage, suggesting coordinated induction of regulatory immunity that may help constrain immune activation and limit immune exhaustion. Together, these results suggest that prior to ATI, the VCs have a “response ready” immune state that engages antiviral and effector programs earlier. These results also support ERRP as a signature that captures intercept-associated transcriptional dynamics.

To examine ERRP expression levels between NCs and VCs, we compared ERRP scores over time. NCs had higher ERRP scores at baseline and at rebound (both p=0.01; **Fig. 4f**). Next, we built piecewise regression models for VCs and NCs relating ERRP score to days from rebound (Methods). We estimated an earlier inflection point in VCs (4.4 days before rebound) than NCs (0.9 days before rebound; **Fig. 4g**), consistent with earlier ERRP upregulation in people with controller phenotypes. Together, these analyses showcase VCs engaging earlier and more functional immune responses than NCs during the intercept.

### Post-intervention controllers have an intermediate immune phenotype

To assess reproducibility of the immune activation observed in REBOUND, we tested whether these patterns could be detected in independent ATI cohorts. We performed GSEA for ERRP enrichment in a previously analyzed, non-interventional ATI cohort of 8 PLWH profiled by single-cell RNA-seq^25^ comparing one baseline timepoint, one pre-rebound timepoint, and one rebound timepoint (**Fig. 5a**). ERRP was strongly enriched in both classical and non-classical monocytes pre-rebound (classical p-adj<1x10^-5^; non-classical p-adj<1x10^-5^) and at rebound (classical p-adj=1x10^-5^, non-classical p-adj=1.9x10^-4^), indicating that ERRP enrichment occurs in distinct cohorts and can be detected with sparse sampling at single-cell resolution.

**Fig. 5.**
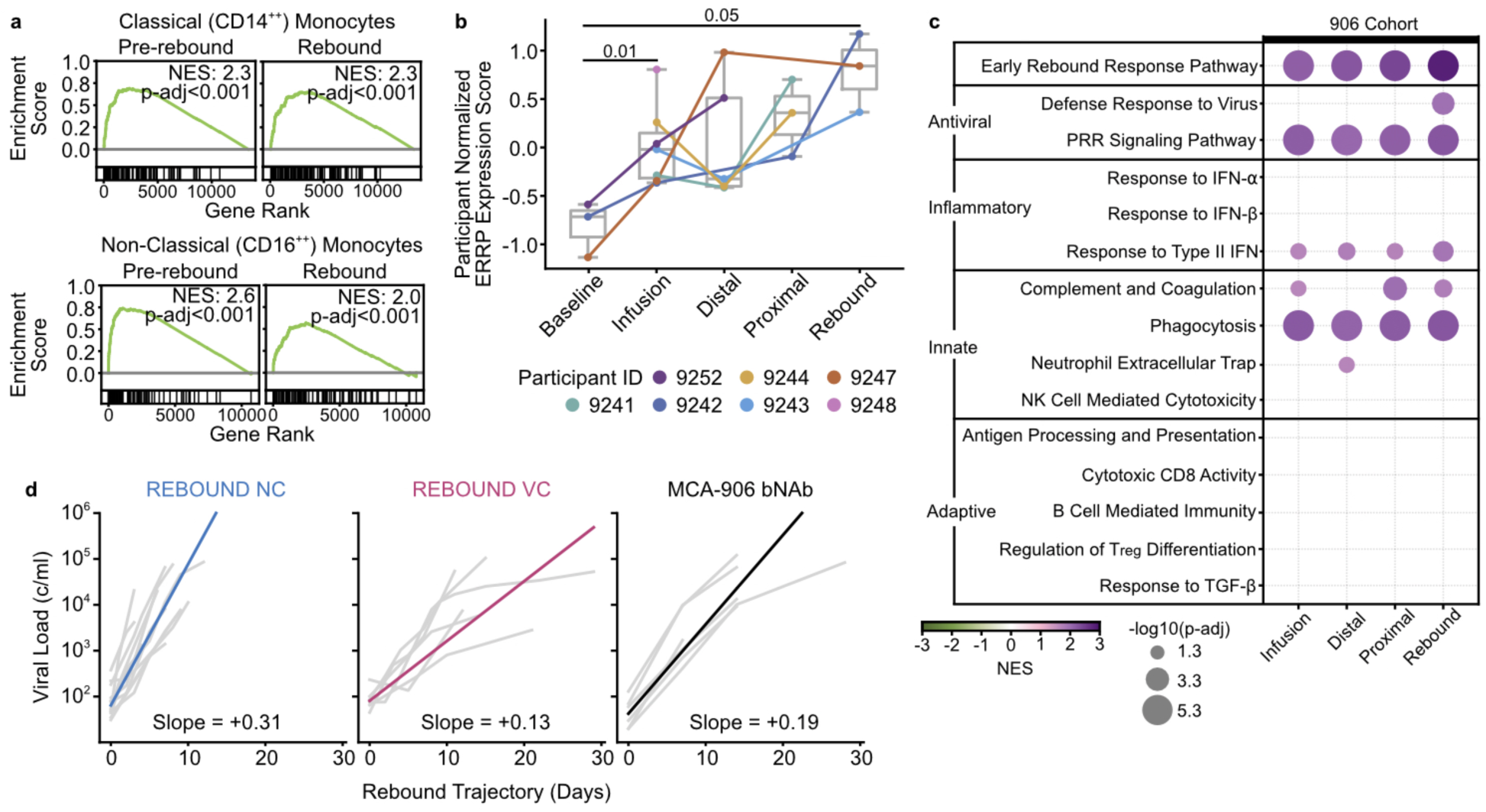
bNAb-mediated viral suppression is associated with early immune activation. **a**, GSEA ERRP enrichment plots in CD14^++^ monocytes (top) and CD16^++^ monocytes (bottom) at pre-rebound and rebound in a second non-interventional ATI cohort. **b**, Average participant-normalized expression of ERRP genes per MCA-906 cohort participant per ATI stage. Points and lines colored by participants. All boxplots represent interquartile range and median across participants. p-values from one-sided Mann-Whitney U test assuming lower baseline expression. **c**, Pathway enrichment across rebound categories in MCA-906 cohort participants. Positive NES indicates enrichment at indicated ATI rebound category; negative NES indicates enrichment at baseline. Full pathway names and reference numbers in Extended Data Table 2. **d**, Rebound slopes of MCA-906 bNAb participants, REBOUND NC, and REBOUND VC overlayed with group median slope. NES=Normalized Enrichment Score.

To test whether ERRP dynamics extend to intervention-delayed rebound, we applied the same analysis to the MCA-906 cohort, in which participants without pre-ART viral control received three infusions of the bNAbs 3BNC117 and 10-1074 at weeks 0, 3, and 6 after ART interruption^4^. We analyzed PBMC transcriptomes from seven participants with bNAb-sensitive reservoirs who exhibited delayed rebound, focusing on five ATI stages: baseline, an early infusion window when bNAb levels were high (weeks 3-9), and distal, proximal, and rebound stages. Participant normalized ERRP expression score increased from baseline to the infusion stage (p=0.01) and to rebound (p=0.05, **Fig. 5b**), and GSEA showed ERRP enrichment across all ATI stages (**Fig. 5c**). Multiple other innate pathways were also enriched prior to rebound, with the strongest signals in ERRP, pattern-recognition receptor signaling, and phagocytosis. We did not observe enrichment of all REBOUND VC-associated effector pathways (including NK cell activity, antigen presentation, or adaptive immune responses, **Fig. 4g**), nor progressive expansion of immune pathway engagement. Consistent with an intermediate phenotype, rebound slope in MCA-906 were slower than REBOUND NC (p=0.0047) but did not differ from REBOUND VC (p=0.31; **Fig. 5d**).

## DISCUSSION

In the REBOUND study, we leverage dense sampling during an observational ATI to understand immune dynamics during the earliest post-ART host-virus intercept and reveal differences in immune activation between individuals with varying control phenotypes. First, we show that viremic controllers rebound later and slower than non-controllers, adding to growing evidence that rebound is not solely a function of reservoir characteristics but is also influenced by contemporaneous host immune pressure^30,43–46^. Second, we observed consistent immune activation before the rebound threshold, including early IFN-⍺1 increases, expansion of non-classical antiviral monocytes, and induction of a conserved early rebound response pathway (ERRP) transcriptional signature. Third, we found that viremic controllers and non-controllers differed in the timing and progressive activation of coordinated innate and adaptive programs, with activation apparent earlier in the controllers. These results support the interpretation that host-mediated control of HIV may require an early, coordinated, pre-rebound immune response.

Viremic controllers exhibit diverse immune signaling during the intercept, including strong enrichment of both innate and adaptive immune functionality. Conversely, non-controllers showed only slight inflammatory activation prior to rebound, with no engagement of functional immune pathways. Controllers also showed greater recruitment of antiviral non-classical monocytes, and increased activation of cytotoxic CD8^+^ T cells. Immediately post-intercept, during the initial viral growth phase, non-controllers showed higher inflammatory activation, while controllers maintained diverse immune functionality. Some of these functions have previously been linked to viral control, but here we describe their activation during the earliest growth-phase of rebound.

In addition to mounting more diverse functional immune responses, controllers also experienced longer periods of immune activation prior to rebound than non-controllers. This is exemplified by ERRP, which is enriched pre-rebound across participants but is activated earlier by controllers than by non-controllers. The pathway includes molecules secreted by infected cells such as MX1, CMPK2, and IFIT1; chemokines that recruit monocytes (CCL2), T cells, and NK cells (CXCL10); and other genes which are induced in response to viral infection. Induction of these genes prior to the rebound threshold suggests that this activation is a direct response to local viral reactivation in tissues. Two explanations could account for earlier ERRP induction in controllers: their more effective response may prolong the interval before systemic rebound, and/or they may initiate ERRP at lower levels of viral activity due to heightened sensitivity of viral sensing. Further studies will be needed to dissect these possibilities or show they work in concordance.

Our data support a model whereby controllers engage diverse immune pathways, potentially facilitated by their lower level of baseline inflammation. We found that non-controllers exhibit higher on-ART inflammation, potentially reflecting differences in chronic immune activation, which has been shown to correlate with latent reservoir activity^47^. Chronic inflammatory signaling can reshape responsiveness to subsequent stimuli and may contribute to an immune system that appears “activated” yet is less capable of mounting a coordinated immune response, perhaps due to upregulation of potent immunosuppressive, counter-regulatory responses^48^. Recent studies also suggest enhanced innate immunity in controllers, which may be mediated by epigenetic programing, and prime the responses of adaptive immunity^49–51^. If this is the case, then controllers, with low baseline inflammation, would have preserved immune responses to low-level tissue HIV reactivation^47^. This hypothesis is supported by studies on type I IFN blockade, which lowers immune activation and has been shown to delay time to rebound^52^ and restore immune function^53^. Based on these results, we propose a model in which controllers sense viral activity early and mount functional immune responses because they have conserved immune coordination. In contrast, non-controllers, who experience higher baseline inflammation, fail to recruit effective functional responses. Interventions pre-ATI and during the immediate post-ART period that seek to shape these early responses will be necessary to untangle how the virus and host interact during this critical period.

Interventional ATI cohorts add to the spectrum of phenotypes in which intercept dynamics can be studied. Here, we studied immune responses in a cohort of ART suppressed individuals who switched to bNAb treatment, where the antiviral activity of these antibodies suppressed viremia for up to 24 weeks. We assume that viral control was largely mediated by direct neutralization of free virus or antibody-mediated clearance of infected cells, however, we observed induction of immune responses during the transition from ART suppression to bNAb treatment. Similar to REBOUND controllers, bNAb recipients increased ERRP expression immediately post-ART cessation and enriched pathways involved in pattern-recognition receptor signaling and phagocytosis, however, they did not enrich expression of adaptive immune responses or NK cell activity. This intermediate profile suggests that low-level virus activity occurs during bNAb therapy, which can be “sensed” immunologically. The degree to which these early immune changes contribute to post-bNAb control, which has been observed in several studies^46,54–57^, remains to be defined. Together, these comparisons motivate strategies that explicitly couple early innate sensing to durable effector responses.

The association between rebound slope and control phenotype has practical implications for future ATI trial design and cure studies. While time to HIV rebound remains a primary endpoint in many studies, it is heavily influenced by the viral threshold used to define rebound, the monitoring cadence, and the participants’ reservoir size^58,59^. Our results suggest that early growth kinetics may better capture the degree to which host immunity constrains expansion of emerging virus and therefore the probability of control. This is particularly important as more early treated individuals take part in cure research: early treated REBOUND participant 211, for example, has a very small reservoir and rebounded much later than other non-controllers, but rebounded with a steep slope indicative of their status as a non-controller. Similar results were observed in another early treated cohort from the Military HIV Research Program^60^. In our bNAb cohort, we observed slopes more similar to REBOUND controllers than non-controllers. Future studies can help determine if this was caused by residual bNAb activity in tissues or if the immune responses generated in conjunction with bNAb treatment were enough to start generating controller-like phenotypes. If validated, slope of rebound, perhaps combined with immune signatures, could provide earlier readouts of induced interventional control without requiring prolonged viremia in ATI participants.

Our study has several limitations. The immune signatures we describe are best understood as direct responses to viral reactivation and replication, likely initiated in tissues and only later becoming detectable in blood. Since our analysis focuses primarily on peripheral blood, we lose spatial resolution and may incompletely capture rare but critical cell populations and tissue-resident responses^23,33,61,62^. Additionally, we show that these phenotypes are directly related to baseline characteristics, which means that our cohort demographics and predominance of subtype B virus directly limit the diversity of baseline states we observed. Even in a relatively homogeneous cohort of largely older men with subtype B, we still found large participant-to-participant variability in baseline measurements and individual trajectories. Larger studies focused on women and individuals from other regions, including those with other viral subtypes, are needed. Additionally, ART restart safety-criteria limited our ability to observe late rebound dynamics in non-controllers, including the possibility of post-treatment control (PTC) in some individuals, although is expected in only 1% of PLWH who initiated ART in chronic infection. Further studies on PTC can elucidate how these individuals might respond during the intercept. Finally, as with all longitudinal human studies, participants experience intercurrent exposures and physiologic variability (e.g., minor infections, vaccinations, other medications), which can introduce noise into immune measurements and contribute to inter-individual heterogeneity.

In the absence of approaches that eliminate the proviral reservoir, a more realistic route to durable HIV-1 remission may be to induce host immunity to control reactivated virus. By defining the timing and composition of immune responses at the host-virus intercept in viremic controller and non-controllers, this work offers a framework to benchmark interventions against a controller-like immune phenotype. A key next step will be to apply this approach to more ATI study participants and to distill the signals identified in REBOUND into smaller, more tractable biomarker panels and to prospectively evaluate whether such markers can simplify ATI monitoring. More broadly, these data highlight how defining the basic immunology of the immune response to rebound in diverse PLWH provides invaluable insights to motivate interventional strategies aimed at delaying rebound and inducing the coordinated aggregate immune response that characterizes viremic control and is likely required for durable ART-free remission.

## METHODS

### Study participants and samples

The Researching Early Biomarkers of Unsuppressed HIV Dynamics (REBOUND) study enrolled twenty people living with HIV (PLWH) on suppressive antiretroviral therapy (ART) who were willing to undergo an observational treatment interruption (NCT04359186). All participants had documented HIV-1, had been on ART for at least 12 months without persistent detectable viremia above 75 copies/ml, and had a screening CD4^+^ T-cell count >350 cells/mm^3^. At enrollment, participants were classified as viremic controllers (VC) if their median pre-ART viral load was <2,000 copies/ml; participants that did not meet these criteria were deemed non-controllers (NC). After ART cessation, participants donated whole blood samples 3 times weekly for the first month of ATI then once weekly until ART restart. In NC, ART was restarted when the viral load measured >200 copies/ml in two subsequent visits. In VC, ART was restarted if the viral load met the following criteria: >50,000 copies/ml for 4 weeks, >10,000 copies/ml for 6 weeks, >2,000 copies/ml for 12 weeks, or >400 copies/ml for 24 weeks. Several VC elected to restart ART before these criteria were met. Study criteria also required ART restart if the CD4^+^ T cell count dropped <350 cells/mm^3^ for two subsequent visits or in the case of acute retroviral syndrome or pregnancy but these events did not occur.

Peripheral blood mononuclear cells (PBMCs) and contemporaneous plasma were isolated from whole blood samples by density centrifugation using Ficoll-Hypaque. Optional additional samples including leukapheresis and lymph node fine needle aspirates (FNA) were taken at baseline and again during the early ATI period; we tried to time the procedures with the initial rebound (predicted to be at or around day 14) whenever possible. Additional tissue samples were taken later in ATI in select participants who did not rebound in 30 days.

### Ethics Statement

The study was approved by the University of California-San Francisco (UCSF) Institutional Review Board and all participants provided written informed consent.

The MCA-906 study (Clinical Trial NCT02825797) ^4^ provided de-identified PBMC samples. The study was approved by the Rockefeller University Institutional Review Board and all participants provided written informed consent.

### Time to rebound and rate of rebound calculations

Rebound is defined as the estimated day a participant’s viral load (VL) reached 200 copies/ml, chosen to align with the lower limit of quantification for globally used clinical viral load assays, and with the World Health Organization (WHO) 2020 threshold of an untransmissible viral load (note: the WHO threshold for viral transmission has since shifted to 1000 copies/ml)^1-3^. To estimate the day of rebound value, a weighted quadratic exponential polynomial model was fit per participant. With log_10_ viral load as *y* and days from ATI day 0 as *x*, this model can be represented as: *log_10_(y) = ax^2^ +bx + c*.

The model was fit with weighted least squares via ‘numpy.polyfit’ function (weights proportional to *√(y)*) using the three samples prior to the first VL above 200 copies/ml and the first sample above 200 copies/ml (and under 1000 copies/ml). Day of rebound was rounded to nearest whole day per participant. These estimates were compared to an alternate method using spline interpolation to confirm robust results. We found no significant differences in estimated day of rebound between the two models (p=0.56, two-sided Wilcoxon test).

Rate of rebound was estimated by calculating the slope between a participant’s first timepoint with a VL >30 copies/ml (limit of detection) and peak viral load (log_10_ transformed) using a least-squares linear regression. Participants who did not reach a minimum viral load of 1000 copies/ml during the ATI were excluded from rate of rebound estimations. To estimate rate for the MCA-906 cohort, a least-squares linear regression was fit between a participant’s first quantified viral load (>20 copies/ml) to peak rebound viral load. If a participant’s first quantified viral load was <200 copies/ml, the last sample at limit of detection was used instead.

### Proviral gag ddPCR

CD4^+^ T cells were isolated from cryopreserved PBMCs with the EasySep Human CD4⁺ T Cell Negative Depletion Kit (StemCell Technologies, #19052). Genomic DNA was extracted using QIAGEN’s DNeasy Blood & Tissue Kit (#69504) or by phenol-chloroform extraction followed by ethanol precipitation. Droplet Digital PCR (ddPCR) was performed to compare the ratio of RPP30 with HIV gag. This ratio was used to calculate the number of gag copies present per million CD4^+^ T cells.

### Lymph node fine needle aspirate (FNA) processing and single-cell library preparation

Baseline FNAs were processed from 12 participants (VC=2, NC=6, Extended Data Fig. 2-3). Lymph node (FNA) specimens were processed immediately after collection. FNA samples were gently triturated and passed through a 70-µm cell strainer. Red blood cells were lysed with ACK lysis buffer then the remaining LNMCs were assessed for cell count and viability. For single-cell analysis, LNMCs were processed using the Chromium GEM-X Single Cell 5′ Reagent Kits v3 platform (10x Genomics) according to the manufacturer’s instructions. 50,000 LNMCs were loaded and processed on Chromium X instrument to generate 100 µl gel beads-in-emulsion (GEMs). After partitioning, GEMs were immediately advanced to reverse transcription. Gene expression and paired V(D)J libraries were generated using the Chromium GEM-X Single Cell 5′ workflow following the manufacturer’s instructions.

### Immune plasma protein abundance measurement via NULISAseq

Immune plasma protein analysis was performed on all participants. Samples were analyzed using Alamar’s Nucleic acid Linked Immuno-Sandwich Assay (NULISA) on the ARGO-HT to detect and measure low level plasma proteins with high sensitivity^63^. NULISA employs a sequential immunocomplex capture and release technique using antibodies conjugated to oligonucleotides. Subsequent analysis of the dsDNA library generated was analyzed by next-generation sequencing (NGS), using the Element AVITI. For quality control, an exogenous reporter was added to each well as an internal control and to normalize data. Pooled plasma was used as a sample and control to assess run suitability, inter-plate and intra-plate CV. Assay buffer was used as a negative control to determine the limit of detection. Sequencing data (FASTQ files) were imported to the Alamar Control Centre (ACC) for downstream processing and QC review. Calculation of NPQ (NULISA protein quantification) values was carried out by normalizing all target reads by the internal control, then dividing those values by target-specific inter-plate control medians, calculated for inter-plate control replicates. Data was rescaled by multiplication with a factor of 10,000 before being log_2_ transformed after the addition of +1 to all values. Participant 203’s samples were flagged during QC for low detectability (<90%) and high internal control reads (>40%). Their NPQ distribution showed a much lower range than rest of the participant samples and was subsequently excluded from downstream NULISAseq analysis as an outlier.

### Flow cytometry

Flow cytometry was performed on 19 participants (VC=6, NC=13, Extended Data Fig. 2-3). Participant PBMC samples from whole blood were randomized, then thawed and stained for two in-tandem high-dimensional flow cytometry panels in 96 well plates using a range of 500,000-2 million cells (panels in Supplementary Data). Cells were resuspended and incubated in 100 µls of Human TruStain FcX™ (Biolegend), MonoBlock (BD), and LIVE/DEAD™ blue (Invitrogen) viability/Fc block stain solution for 15 minutes at RT. Cells were washed in FACS wash buffer (2% FBS) and stained in 50 µls of panel-specific tittered antibody cocktails (Supplemental Methods) in BD Brilliant Stain Buffer for 30 minutes at 4°C. Cells were washed twice and fixed in 100 µls of 4% PFA for 20 minutes, then washed and stored in 200 µls in 96 wells plates at 4C in preparation for flow cytometry analysis in the subsequent 48 hours. Single stain compensation controls and Fluorescence Minus One (FMO) were stained using the same method. FMOs were used to standardize gating between batches for each experiment. All flow cytometry analysis was performed using the same FACSymphony A5 instrument (special order product, BD Biosciences, San Jose, CA). FMOs and samples were acquired via High Throughput Screening device in 96 well plates. Subsequent data was analyzed using FlowJo v10.10 software (gating in Extended Data Fig. 7-8).

### Cytometry by Time of Flight (CyTOF)

CyTOF analysis was performed on 15 participants (VC=6, NC=9, Extended Data Fig. 2-3), as described previously^30^. 2–4 million cells were stained for mass cytometry (panel in Supplemental Methods). PBMCs were thawed and only samples with over 70% viability were used for analysis. Dead cells were marked by incubating the samples for one minute with 25 mM cisplatin (Sigma-Aldrich) in phosphate buffered saline (PBS) with EDTA, performed surface staining with metal-tagged antibodies in PBS with 0.5% bovine serum albumin (BSA) for 30 min at room temperature, fixed and permeabilized cells following manufacturer’s instructions for the eBioscience Foxp3/Transcription Factor Staining Buffer Set (Thermo Fisher Scientific), barcoded samples using mass-tag cellular barcoding reagents diluted in Maxpar Barcode Perm Buffer (Standard BioTools, South San Francisco, CA, USA), combined up to twenty barcoded samples into a single tube, performed intracellular staining with antibodies diluted in eBioscience Foxp3/Transcription Factor kit perm wash (Thermo Fisher Scientific), fixed cells in freshly prepared 2% paraformaldehyde (Electron Microscopy Sciences, Hatfield, PA, USA) in the presence of a DNA intercalator, and then washed and ran cells on the Standard BioTools CyTOF 2 mass Cytometer within one week of staining.

After data acquisition, normalization and debarcoding of samples was performed using Premessa (v.0.3.4). FCS files were imported into CellEngine (CellCarta). All manually gated live, CD45^+^ single cells were downloaded as .fcs files from CellEngine and first batch corrected using cyCombine (v.0.3.0). Batch-corrected files were re-uploaded to CellEngine and gated on total CD45^+^ cells. Traditional hierarchical gating (Extended Data Fig. 9 for gating strategy) was applied as previously described^30^ to identify standard immune populations.

### Bulk RNA-seq

RNA-seq was performed on 19 participants (VC=6, NC=13, Extended Data Fig. 2-3). Participant samples were randomized and PBMCs from whole blood were placed in RLT buffer + 40 mM DTT. RNA was extracted and purified from cell lysate using the Qiagen RNeasy Micro Kit according to the manufacturer’s protocol. RNA quantity and quality was assessed using the Qubit high sensitivity RNA assay and the Agilent Tapestation 4200 High sensitivity RNA assay. Using a normalized input of 50ng from each sample, the KAPA mRNA Hyperprep Kit was used according to the manufacturer’s protocol to enrich for mRNA and create RNA-seq libraries for next generation Illumina sequencing. Library prepped samples were quantified and verified for appropriate size using the Qubit 1x dsDNA assay and the Agilent Tapestation D1000 assay. Libraries were normalized to the same concentration and pooled for sequencing. The pooled concentration was verified using the Qubit 1x dsDNA assay prior to sequencing using the Illumina Novaseq X instrument.

### RNA-seq count quantification and preprocessing

RNA-seq count quantification was performed on raw FASTQ files using Salmon v1.10.1 (using Ensembl Version 112 and Homo sapiens GRCh38 to build the Salmon index) for all bulk RNA-seq datasets. Genes with low expression were removed: in the REBOUND cohort this included all genes with fewer than 10 counts across minimum 10 samples; in the 906-bNAb cohort genes with fewer than 5 counts across minimum 10 samples were removed. For the lymph-node scRNA-seq data, pseudobulked celltype data genes with fewer than 2 counts within at least 5 samples were filtered out. Downstream analysis was performed in Python (v3.11.5). Differential gene expression used DESeq2 (PyDESeq2; v0.4.10). Batch numbers were included in design factors for datasets run in multiple batches. Longitudinal contrasts include participant-IDs as design factors for paired analysis. Adjusted p-values represent Benjamini-Hochberg (FDR-BH) corrected p-values for multiple hypothesis testing. Generally, a |log_2_(FC)|>0.5 and p-adj<0.05 cut-off was used to define differentially expressed genes across all contrasts. For visualization purposes, volcano plot’s x-axes were limited between -4 and 4 with complete lists of DEGs in Supplementary Table 3.

### ERRP gene pathway definition

To robustly identify genes consistently upregulated from baseline to rebound, we performed two batches of REBOUND bulk RNA-seq. Batch 1 contained 93 samples from a subset of the REBOUND-cohort participants (5 VC, 8 NC) and was used to find an initial signature of 140 differentially expressed genes between baseline and rebound (|log_2_(FC)|>0.5, p-adj<0.05). Red blood cell associated genes (HBB, HBA1/2, and ALAS2) were removed from the analysis. Batch 2 contained an additional 122 samples from the REBOUND-cohort (70 samples from participants in batch 1, 39 samples from 5 new NC, 13 samples from 1 new VC). Batch 2 samples were combined with the initial batch, and the 138 genes were re-tested for differential gene expression. 116 genes remained significant (|log2 fold change|>0.5, p-adj< 0.05). These 116 genes were treated as a pathway (Early Rebound Response Pathway, ERRP) for downstream gene-set enrichment analysis. 8 genes of the 116 genes in ERRP are expressed higher in baseline than rebound.

### GSEA Pathway analysis

Gene set enrichment analysis (GSEA) was run through GSEApy (v1.1.3) with gene libraries KEGG 2021^64^, GO:BP 2023^65^, and additional CD8 pathways alongside ERRP. Genes were ordered prior to GSEA using a metric rank = sign(log_2_FC) × -log_10_(p-value). Enrichment scores (ES) were estimated using the GSEApy ‘prerank’ algorithm with N=1000 permutations. Normalized enrichment scores (NES) were used to summarize pathway activity and pathways with FDR q-value<0.05 threshold were considered significantly enriched.

### Single-cell RNA and TCR sequencing count quantification, preprocessing, and quality control

scRNA and TCRseq count quantification was performed on raw FASTQ files using CellRanger (v7.1.0) using the *Homo sapiens* reference GRCh38 for both GEX and VDJ libraries. Next, ambient RNA was removed using SoupX (v1.6.2) and the adjusted counts were rounded to integer values. Then, doublets were removed using DoubletFinder (v2.0.3), scrublet (v0.2.3), and cells with two beta chains were removed. Next, low-quality cells were removed by applying thresholds of cell quality metrics separately for each library. Across all libraries, retained cells had 1,500-15,000 UMIs, 660+ genes, 10-55% ribosomal UMIs, 0-15% mitochondrial UMIs, 0-3% hemoglobin UMIs, 0-0.4% platelet UMIs, and 0-20% of UMIs from the gene with the most UMIs.

### Single-cell RNA-seq sample integration and clustering

Cells were clustered using Seurat (v5.0.0), where cells from each batch were encoded as a layer in the Seurat object and then normalized using SCTransform with ‘variable.features.n’ set to 3,500. Next, to prevent unwanted sources of variation from driving clusters, genes from the following categories were removed from Highly Variable Genes (HVG): variable TCR, variable BCR, X chromosome, Y chromosome, ribosomal, mitochondrial, hemoglobin, and platelet genes. After pruning the HVG, the genes with the lowest dispersion were removed until only 3,000 HVG remained. These HVG and the normalized data were used for the remainder of the clustering pipeline, including, PCA calculation using 50 dimensions and batch normalization performed with IntegrateLayers with ‘method’ set to ‘scVIIntegration’, ‘normalization.method’ set to ‘SCT’, ‘ndims’ set to 50, and ‘max_epochs set’ to 100. After batch normalization, clustering was performed with RunUMAP and FindNeighbors using 50 scvi dimensions, followed by FindClusters with a resolution of 0.7. Cell type labels were assigned by using a combination of Azimuth annotations and marker gene profiling per cluster (Extended Data Fig. 10).

### Expression scores

Expression scores for the ERRP and protein set were calculated by averaging normalized gene/protein expression within pre-defined gene/protein sets. Each set was defined by differential expression/abundance analysis between baseline and rebound so expected direction of change for each gene/protein was determined by the direction of change in that analysis. Therefore, within each set, genes/proteins expected to be downregulated had their expression values sign inverted (multiplied by -1), whereas genes/proteins with expected upregulation contributed to expression values unchanged. Expression score was then defined as the mean of all values per sample for a given gene/proteins set. Thus, expression scores represent a single aggregate activity measurement per set per sample. To compare trends in expression/abundance over time between participants, we calculated a participant normalized expression score. In this case, before averaging, values were normalized within-participant by z-scoring each value relative to that participant’s own distribution: the participant-specific mean was subtracted from each value and divided by the participant-specific standard deviation, yielding standardized values with mean = 0 and unit variance within participant.

### Continuous modeling

We used standard python modules for modeling expression scores. To assess if an expression score could predict how far a given timepoint was from rebound, we used Pymer4 (v0.8.2) and Statsmodel (mixedlm, v0.14.2) for mixed-effects modeling, with all predictors entered as fixed effects and participant ID specified as a random intercept to model between-subject baseline variability. Conditional R^2^ was used to measure model goodness-of-fit which accounts variance explained for by both fixed and random effects of the model^66^. To capture the initial rebound trajectory, timepoints for mixed effects modeling were bounded from 40 days prior and 10 days post rebound.

Breakpoint modeling was calculated using pwlf (v2.5.1) python library specifying 2 segments without pre-determining breakpoint. The package identifies breakpoint locations using optimizations to minimize the sum of squares error.

### Statistical Analysis

Statistical analyses were primarily performed in Python using standard packages Scikit-learn (v1.5.1), SciPy (v1.14.0), and Statsmodel (v0.14.2). Additionally standard data processing packages such as Numpy (v1.26.4) and Pandas (v2.2.2) were used in analysis. Mann-Whitney U tests and Wilcoxon tests used the median value per participant per rebound stage to handle unequal number of samples. Adjusted p-values represent Benjamini-Hochberg (FDR-BH) corrected p-values for multiple hypothesis testing.

## Acknowledgements

The REBOUND study was funded the Gates Foundation (INV-002707 to L.B.C.). The SCOPE cohort was supported by the UCSF CFAR (P30 AI027763), the CFAR Network of Integrated Clinical Systems (R24 AI067039) and the Delaney AIDS Research Enterprise (DARE; UM1AI164560). Additional funding support was provided by the Gates Foundation (INV-055706 to A.K.S.), the NIH (UM1 AI164560, UM1 AI164565, U54 AI170792, to L.B.C., R01 AI170239 to R.L.R., K23 AI157875 to M.J.P University of Washington Center for STD and AIDS Training Grant [T32AI07140 to A.F.S.]), and the Pew Research Scholars Program (to L.B.C.). This research was supported by NIH P30 CA015704 of the Fred Hutch/University of Washington/Seattle Children’s Cancer Consortium, which includes the Flow Cytometry and Genomics & Bioinformatics Shared Resource. In addition, research reported in this publication was supported by the UCSF-Bay Area Center for AIDS Research, an NIH-funded program under award number P30AI027763 which is supported by the following NIH Institutes and Centers: NIAID, NCI, NIDA, NICHD, NHLBI, NIA, NIMHD, NINR, NIDDK, NIDCR. This project was supported by the National Center for Advancing Translational Sciences, National Institutes of Health, through UCSF-CTSI Grant Number UL1 TR001872. Its contents are solely the responsibility of the authors and do not necessarily represent the official views of the NIH. We are grateful to the primary care and HIV providers for the study volunteers, who communicated with the research team to coordinate in preparation for and management of the ATI and ART restart. We thank Michel C. Nussenzweig for generously providing samples used in this work; Julia Jackson, formerly in the Cohn Lab, for foundational technical contributions; the UCSF SCOPE team, especially Emily Fehrman and Lynn Ngo, for clinical coordination; Monika Deswal and Elnaz Eilkhani for regulatory support; the UCSF Core Immunology Laboratory and AIDS Specimen Bank for biospecimen processing, storage, and distribution; Jeffrey Martin and Melissa Krone for database support; Tze Wai Tiffany Shing and Dominic Lung from the UCSF Department of Pathology for assistance in coordinating the lymph node FNAs; all current and former members of the Cohn and Lehman Labs for helpful discussion; Thuy Dao and Monica Lopez-Bernal for research and administrative support; Mohit Jain for the support and coordination of experiments provided by Sapient; Alexander Aravkin, and Aaron Hudson for analytical support; Daniel Reeves, David Sherman, and Andrew McGuire for critical reading of the manuscript; and members of the Gates HIVRC consortium, especially Joseph (Mike) McCune and Heather Ann Brauer. Finally, we would like to thank the participants of all the studies presented here. Their commitment to HIV research makes studies like these possible.

## Author Contributions

A.F.S., W.A. and N.dlF. contributed equally to the work. The project was jointly supervised by S.G.D. and L.B.C.. M.J.P., R.H., S.G.D. and L.B.C. designed the clinical trial. M.J.P, S.G.D., R.H., A.E.R., T.F., M.C.W., V.T. conducted the clinical study. M.J.P. and T.D. analyzed the clinical data. P.S., C.D.l.S.V., P.V., C.G., and S.L. performed the lymph node fine needle aspirate collections. N.dlF. performed bulk RNA-seq and flow cytometry experiments. A.F.S. led experimental coordination for the plasma protein abundance experiments and, together with E.W., measured HIV DNA. D.A.S. and R.L.R. performed the CyTOF analysis. D.A.S., G.G. and A.B.A. performed and G.F.K. and R.L.R. oversaw FNA single-cell RNA-seq pre-processing. W.A. performed all other data pre-processing, and conducted the computational and statistical analysis with the support of A.F.S. and input from B.Z. and A.K.S.. M.C. provided samples from the MCA-906 study. The manuscript was written by A.F.S., L.B.C, and S.G.D. with support from W.A. and the rest of the team. All authors edited and approved the final manuscript

## Competing interests

M.J.P. serves on a DSMB for American Gene Technologies. S.G.D. reports consulting fees from AbbVie, GSK, Hookipa, American Gene Technologies and Immunocore; owns Tendel stock; and receives research support from Gilead. M.C. served on a Gilead scientific advisory board. A.K.S. reports compensation for consulting and/or SAB membership from Honeycomb Biotechnologies, Cellarity, JnJ, Ochre Bio, Danaher, Parabilis Medicines, Passkey Therapeutics, Conquest Technologies, Relation Therapeutics, IntrECate Biotherapeutics, and Dahlia Biosciences unrelated to this work.

## Data Availability

All data will be made publicly available upon publication.

## Materials & Correspondence

Requests should be addressed to Lillian B. Cohn or Steve G. Deeks.

---

**Extended Data Fig. 1:**
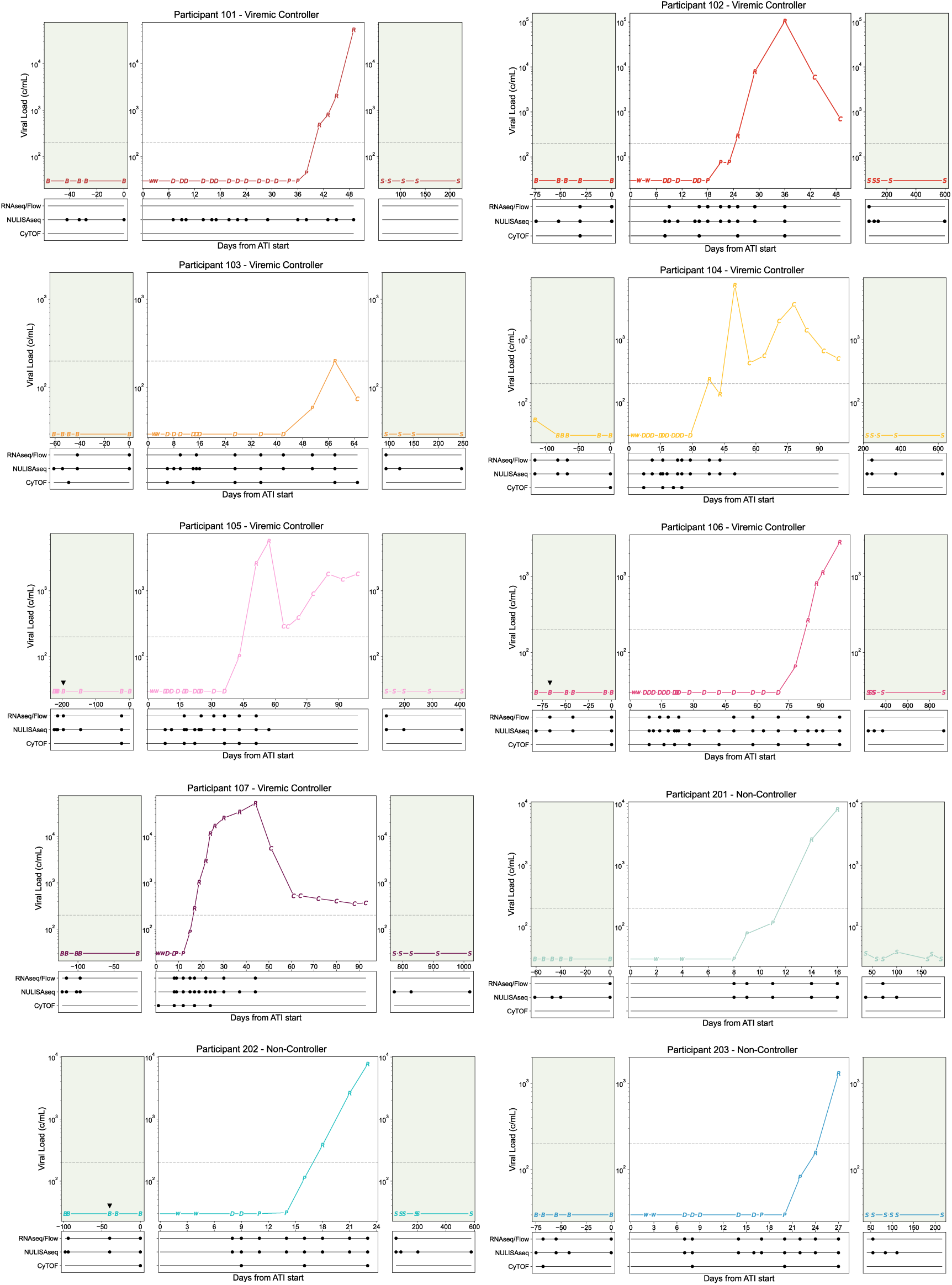
Viral load and sampling from Participants 101-203. Viral load curves by participant, points represent ATI stage: B=Baseline, W=ART washout, D=Distal, P=Proximal, R=Rebound, C=Rebound-Control (samples not used in analysis), S=Follow-up. Dotted grey line at rebound threshold of 200 copies/ml, green background shading indicates ART. Samples used in analysis are annotated below the viral load plot. Black triangle (▾) above viral load point represents baseline FNA.

**Extended Data Fig. 2:**
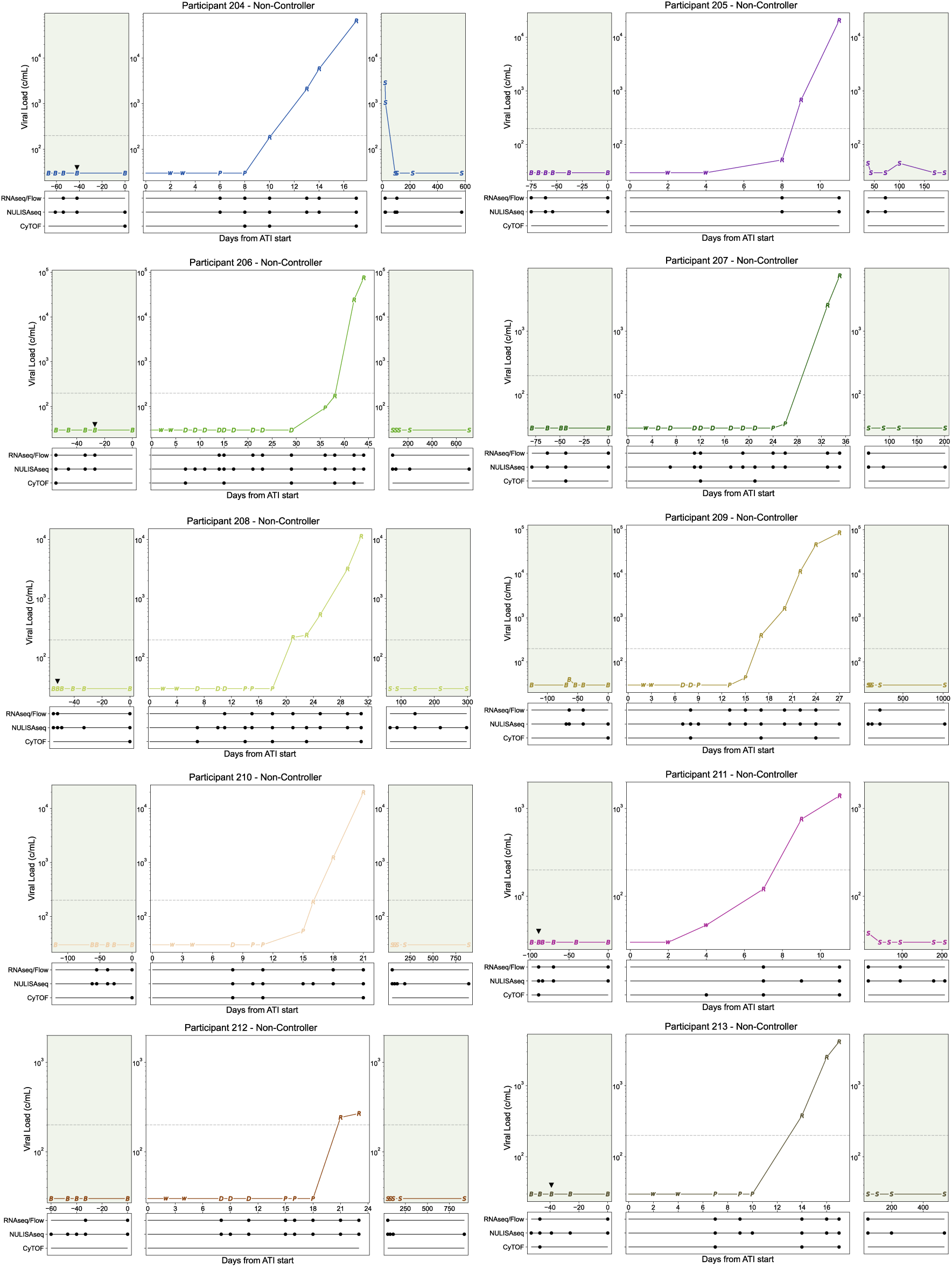
Viral load and sampling from Participants 204-203. Viral load curves by participant, points represent ATI stage: B=Baseline, W=ART washout, D=Distal, P=Proximal, R=Rebound, C=Rebound-Control (samples not used in analysis), S=Follow-up. Dotted grey line at rebound threshold of 200 copies/ml, green background shading indicates ART. Samples used in analysis are annotated below the viral load plot. Black triangle (▾) above viral load point represents baseline FNA.

**Extended Data Fig 3:**
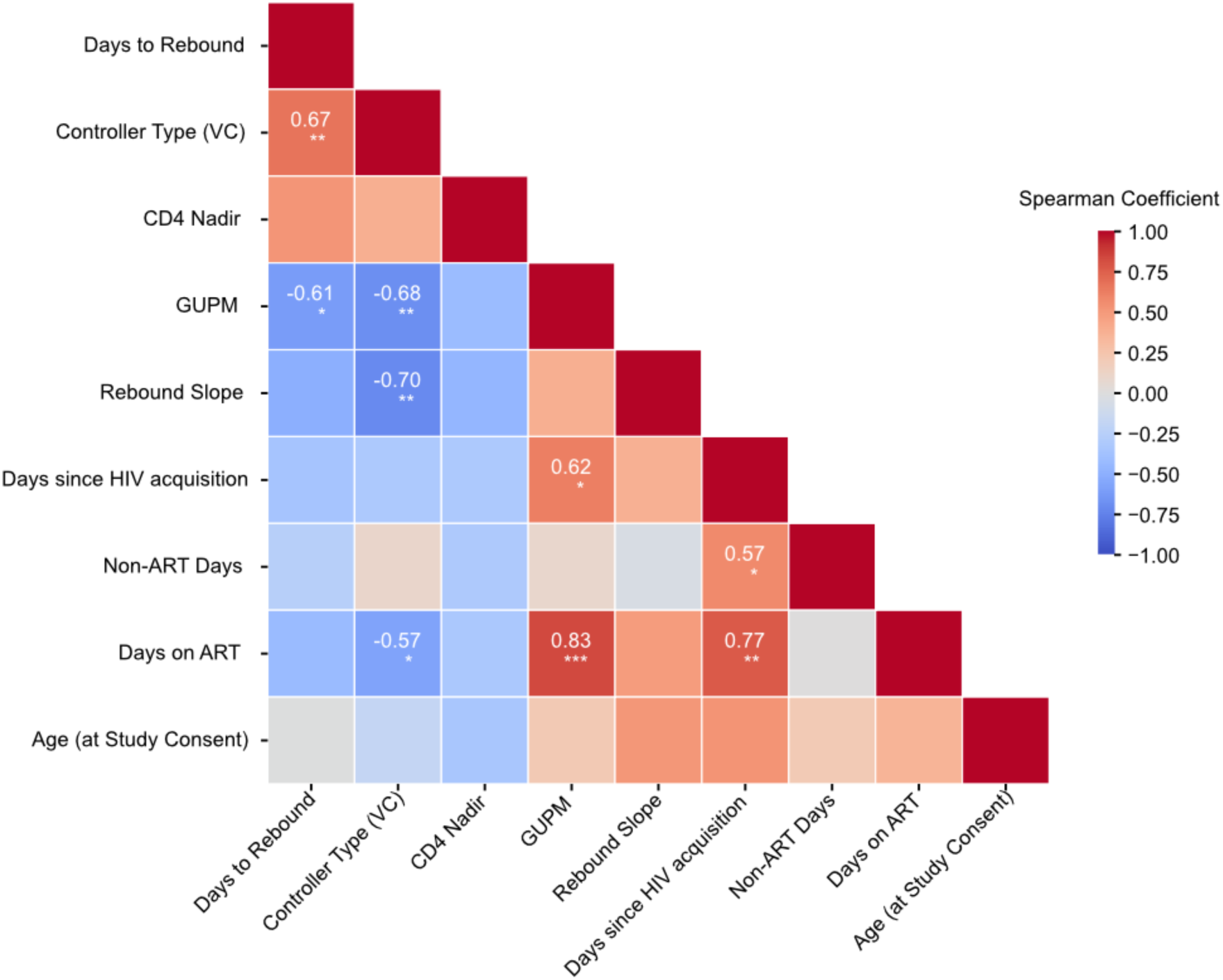
Correlation between participant characteristics. Heatmap of Spearman correlation coefficients between participant characteristics. Color indicates correlation coefficient; significant coefficients are displayed on corresponding square. * FDR<0.05, ** FDR≤0.01, *** FDR≤0.001

**Extended Data Fig. 4:**
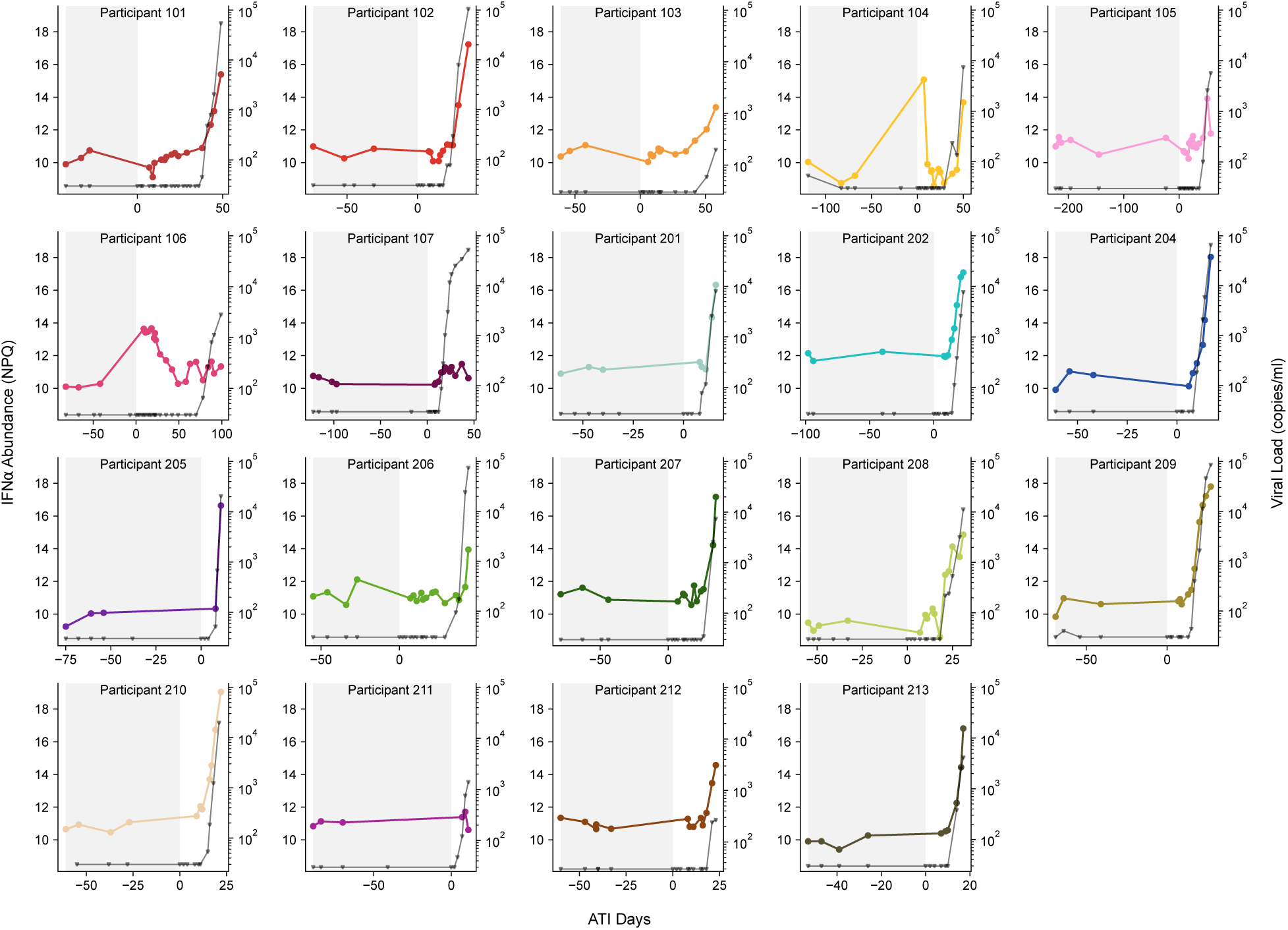
IFN-⍺1 abundance association with viral load in REBOUND participants. Viral load curves for each REBOUND participant (grey lines, triangles, right axis) overlayed with IFN-⍺1 abundance (colored lines, circles, left axis). Grey shading indicates ART. ATI day 0 indicates the first day of the ATI.

**Extended Data Fig. 5:**
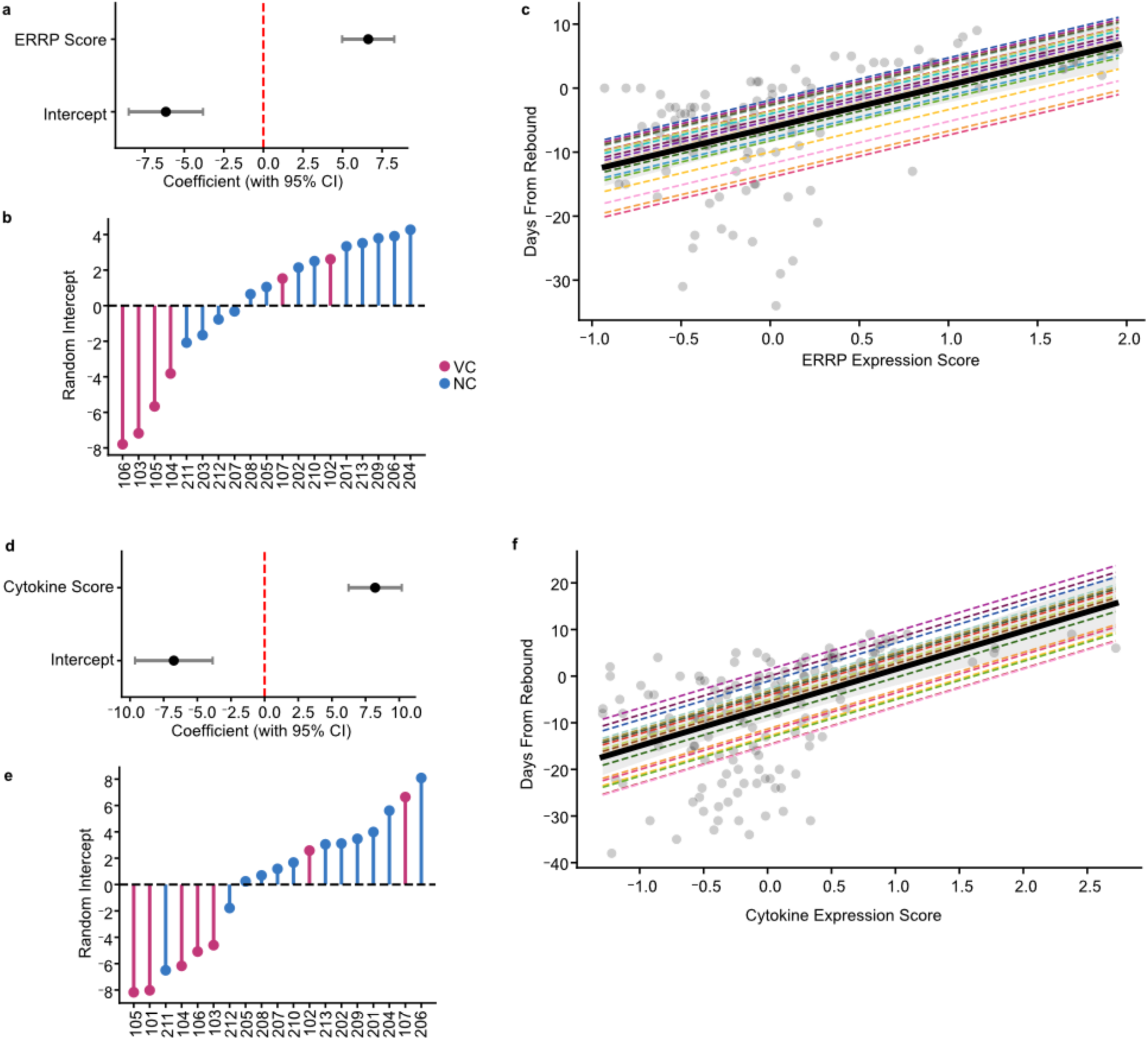
Participant-specific linear mixed effects models associated with time to rebound. Model characteristics of participant-specific linear mixed effects models associating time to rebound with ERRP/Protein score (**a**,**b**,**c**) and soluble protein score (**d**,**e**,**f**). (**a**,**d**) Forest plots showing the fixed-effect coefficients for the measured variable and intercept with 95% confidence intervals. (**b,e**) Lollipop plots showing the distribution of random intercepts across participants used in modelling, reflecting inter-participant variability. Lines colored by participant’s controller type. (**c,f**) Model predicted relationship between prediction (ERRP Score/Protein Score, x-axis) and response (Days from Rebound, y-axis). Gray points represent observed data, thick black line indicates population-level fixed effects. Thin multi-colored dashed lines show participant-level predictions with shared slopes and varying intercepts.

**Extended Data Fig. 6:**
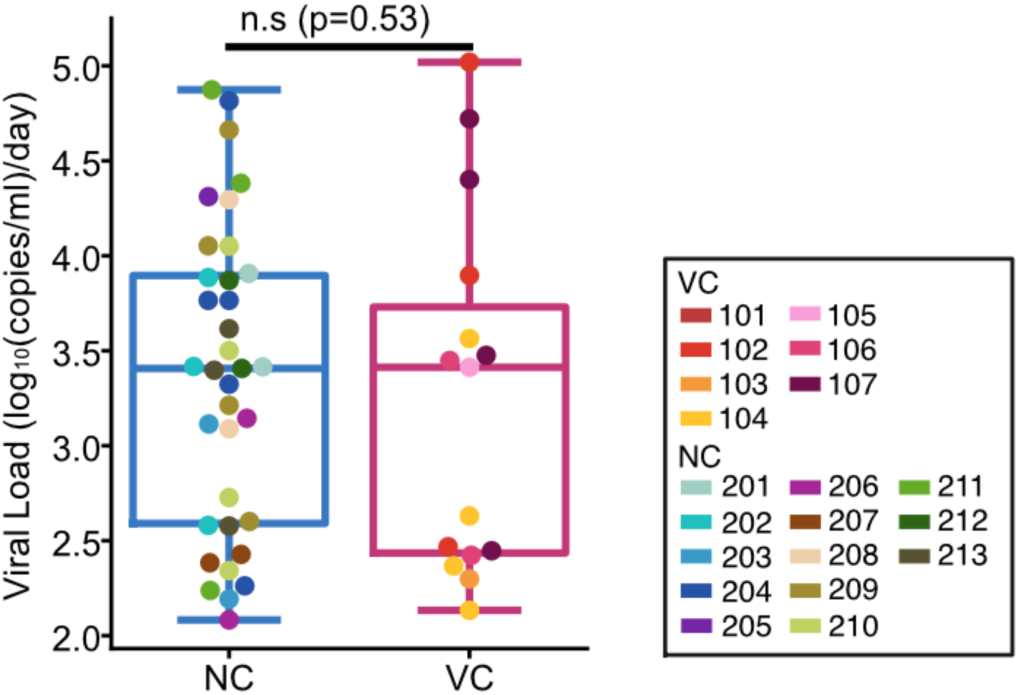
Viral loads of rebound stage timepoints between VC and NC. Viral loads of rebound stage timepoints used for analysis compared between VC (left) and NC (right). p-value from two-sided Mann-Whitney U test.

**Extended Data Fig. 7:**
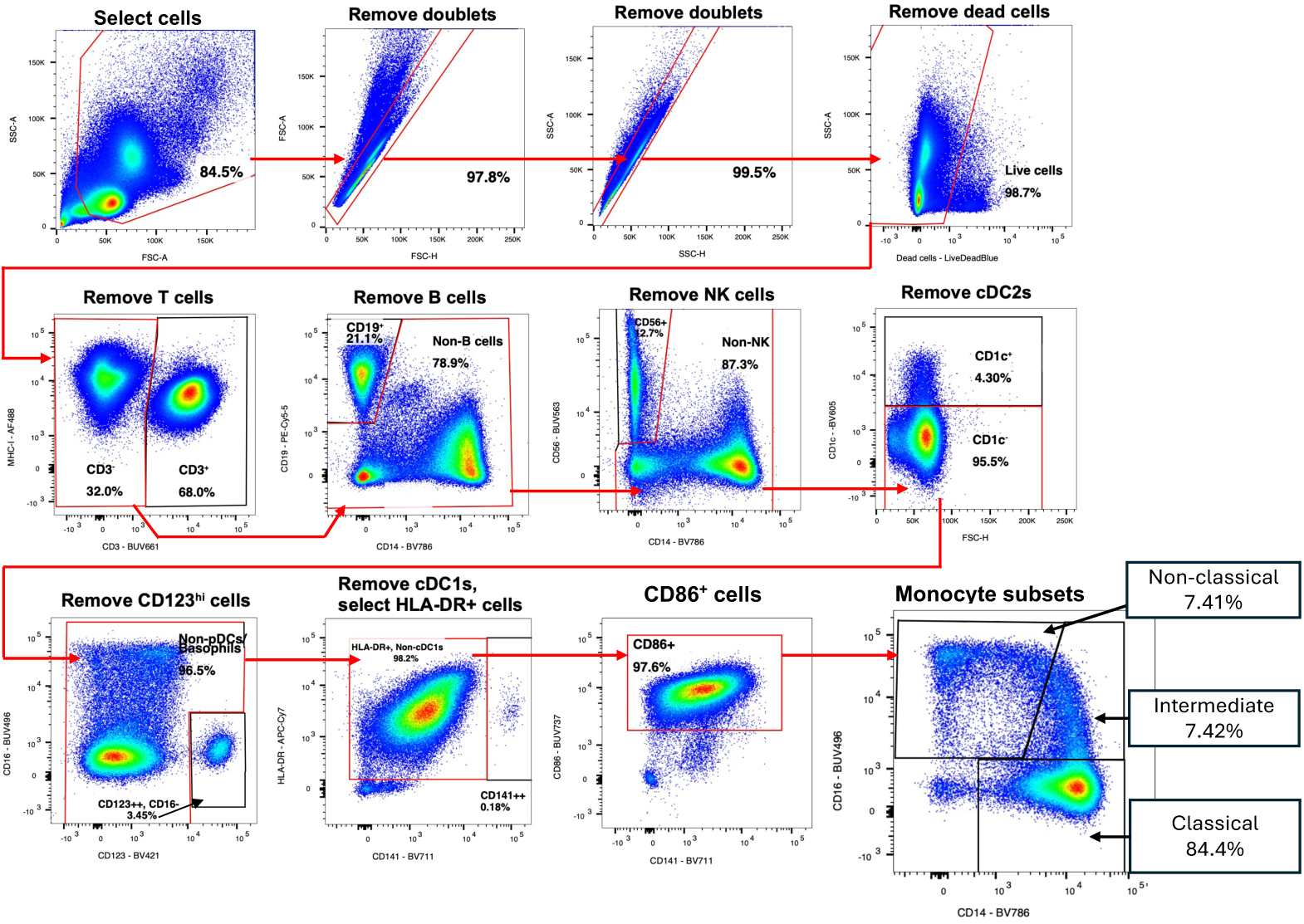
Flow Cytometry gating strategy for myeloid cell panel.

**Extended Data Fig. 8:**
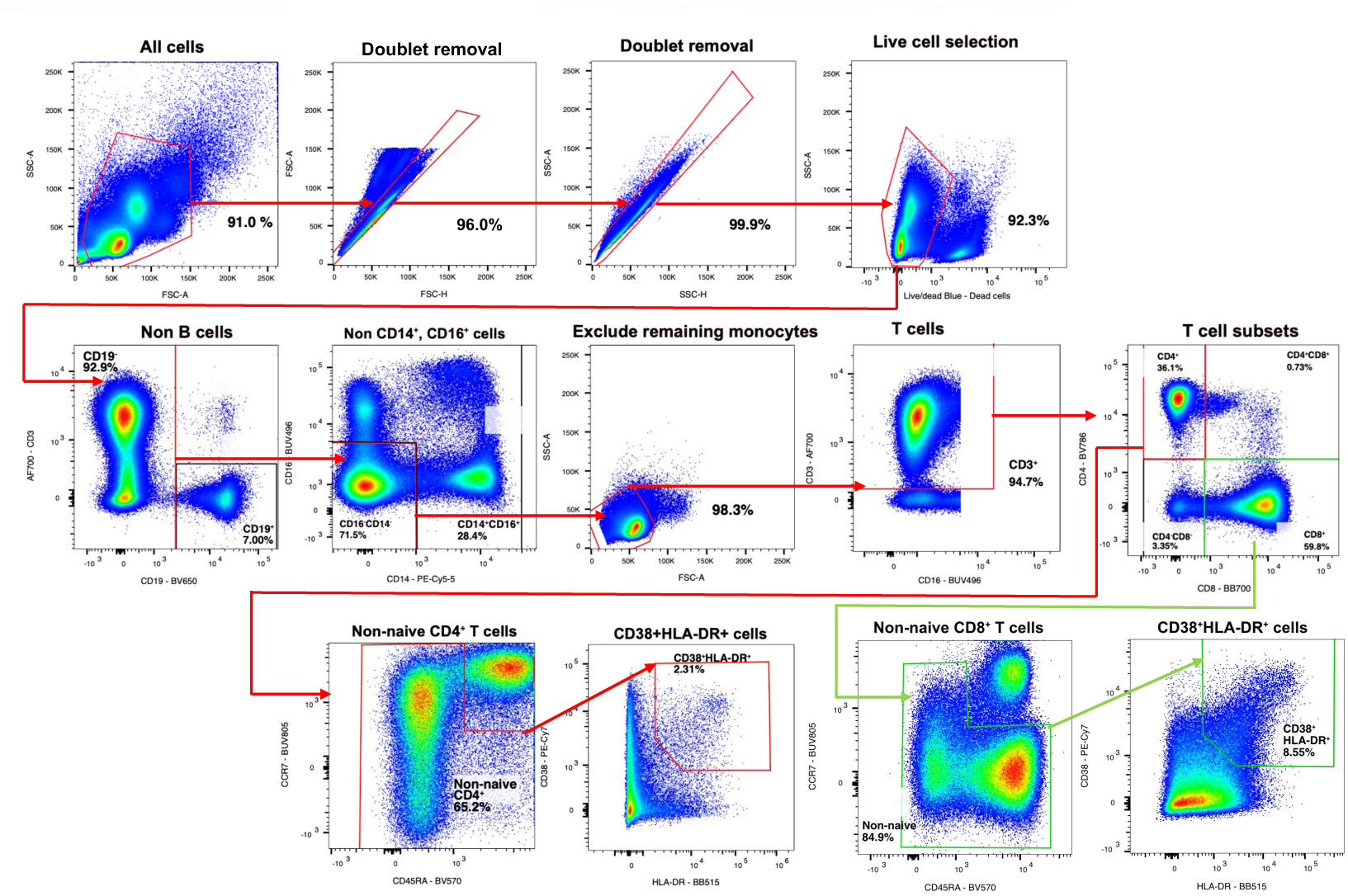
Flow Cytometry gating strategy for T cell panel.

**Extended Data Fig. 9:**
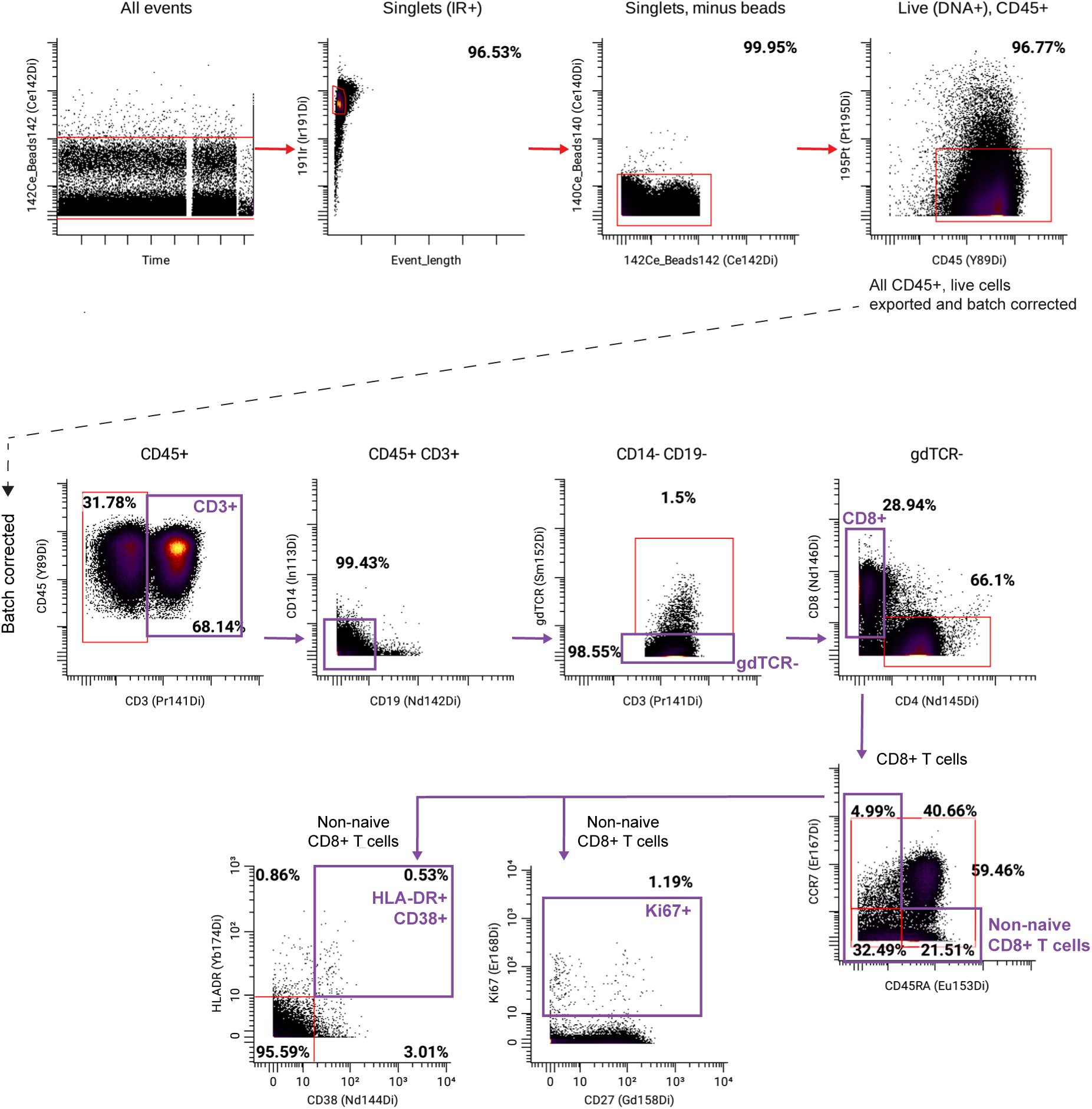
CyTOF gating strategy for T cell panel.

**Extended Data Fig. 10:**
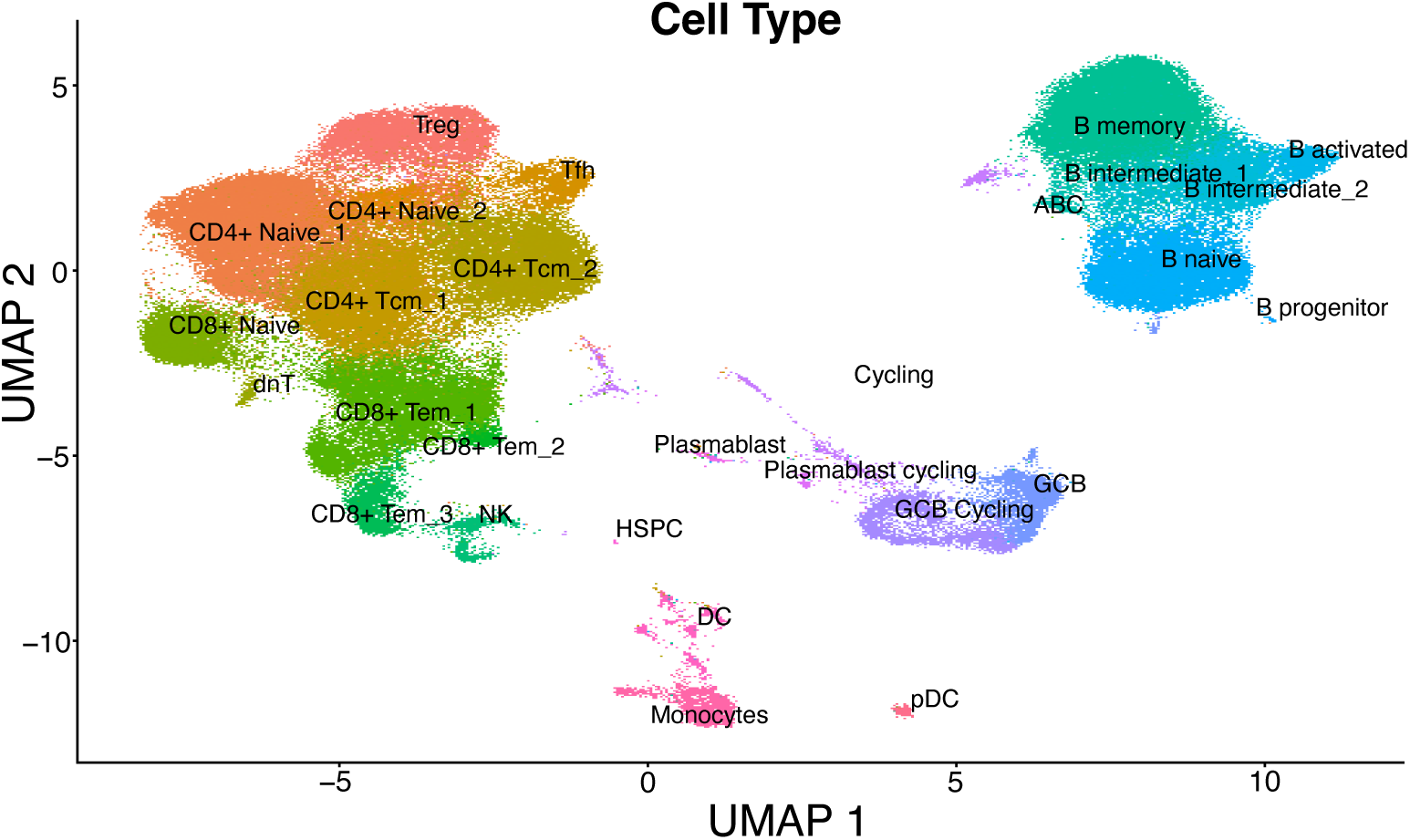
Single cell UMAP of FNA samples.

**Extended Data Table 1.** REBOUND Cohort Participants.

**REBOUND Cohort Participants**
| **Variable** | **Overall** | **Non-Controllers** | **Viremic Controllers** |
| --- | --- | --- | --- |
|  | N = 20 | N = 13 | N = 7 |
| **Age at Study Consent (years)** | 59.5 (32–75) | 63 (41–75) | 56 (32–69) |
| **Gender** |  |  |  |
| Cis man | 15 (75%) | 9 (69%) | 6 (86%) |
| Cis woman | 2 (10%) | 1 (8%) | 1 (14%) |
| Trans woman | 2 (10%) | 2 (16%) | - |
| Non-binary | 1 (5%) | 1 (8%) | - |
| **Race** |  |  |  |
| White | 14 (70%) | 8 (62%) | 6 (86%) |
| African American or Black | 3 (15%) | 2 (15%) | 1 (14%) |
| Mixed Race | 1 (5%) | 1 (8%) | - |
| Unknown/Not Reported/Declined to Answer | 2 (10%) | 2 (15%) | - |
| **Ethnicity** |  |  |  |
| Hispanic or Latino | 3 (15%) | 3 (23%) | - |
| **Viral Subtype** |  |  |  |
| Subtype B | 19 (95%) | 13 (100%) | 6 (86%) |
| Subtype AE | 1 (5%) | - | 1 (14%) |
| **Years from HIV acquisition to ART start** | 4.9 | 5.0 (0.02-16.7) | 4.8 (1.7-23.6) |
| **Years on ART prior to ATI** | 13.5 | 18.9 (4.9-26.7) | 8.2 (1.6-17.0) |
| **CD4<sup>+</sup> T Cell Nadir** | 294.5 | 211<br>(29–836) | 400<br>(249–1114) |
| **Median pre-ART Viral Load (copies/ml)** |  | 8624<br>(120-27000) * | 102<br>(ND-1742) |
| **GUPM (Gag units per million CD4<sup>+</sup> T cells)** | 181.4 | 472.6 (0.0–1124.5) | 31.4 (0.0–178.7) |
| **Rebound Slope (log<sub>10</sub>(copies/ml) per day)** | 0.27 | 0.31 (0.18–0.85) | 0.13 (0.08–0.27) |
| **Days to Rebound (200 copies/ml)** | 21.5 | 17 (7–38) | 40 (17–83) |

Results displayed as: median (minimum-maximum)
ND = Not Detected
\* Median pre-ART viral load not known for 6 NC participants, NC with median pre-ART VL=120 was acutely treated and excluded from median calculation

**Extended Data Table 2:**
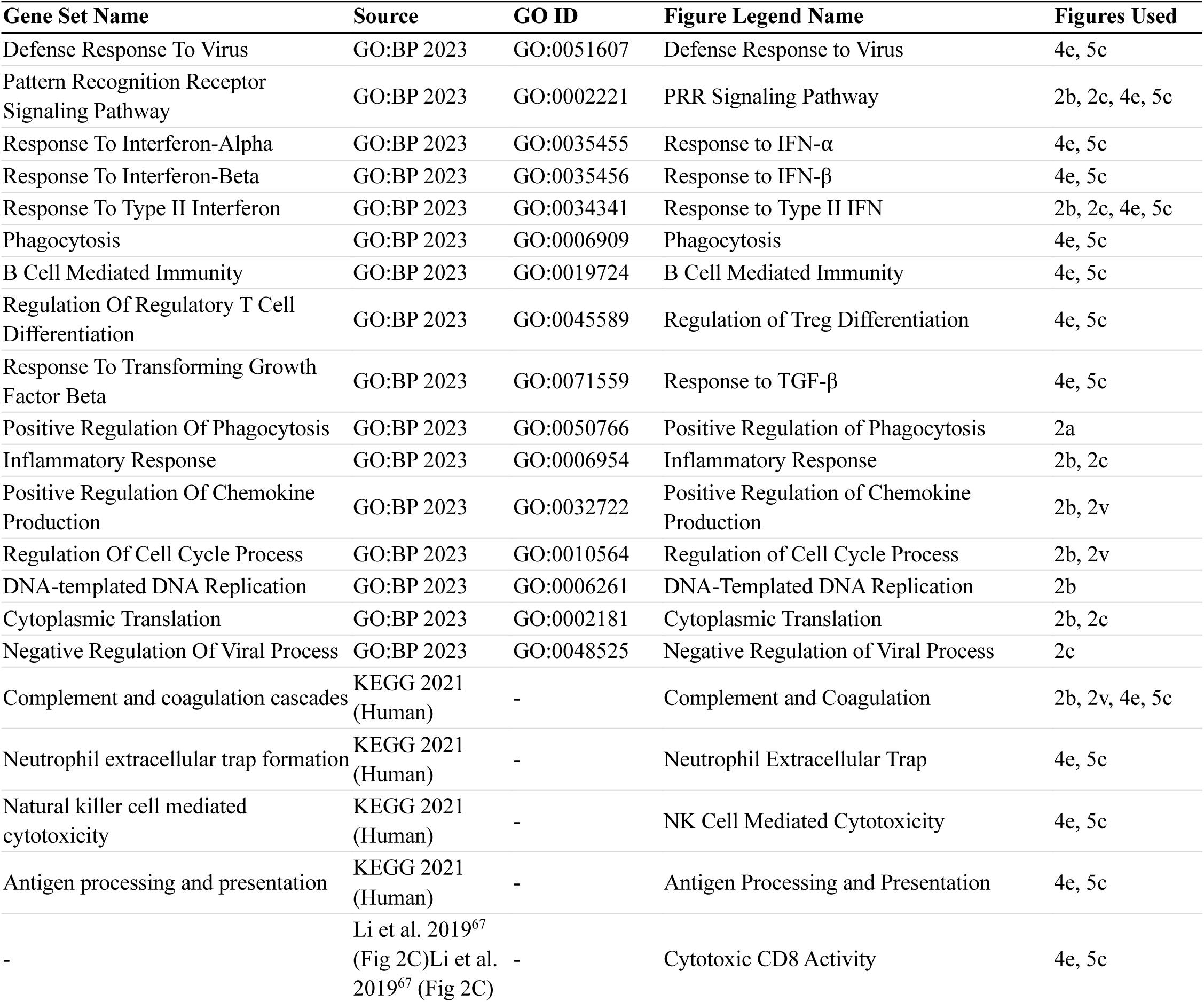
Gene Set Terms.

